# Predicting systemic and pulmonary tissue barrier concentration of orally inhaled drug products

**DOI:** 10.1101/2022.03.10.483633

**Authors:** Narender Singh, Ravi Kannan, Ryan Arey, Ross Walenga, Andrew Babiskin, Andrzej Przekwas

## Abstract

The complex physiology and anatomy of the lungs and the range of processes involved in pulmonary drug transport and disposition make it challenging to predict the fate of orally inhaled drugs. This study aimed to develop an integrated computational pharmacology approach to mechanistically describe the spatio-temporal dynamics of inhaled drugs in both systemic circulation and site-specific lung tissue. The model included all the physiologically relevant pulmonary processes, such as deposition, dissolution, transport across lung barriers, and mucociliary clearance, to predict the inhaled drug pharmacokinetics. For validation test cases, the model predicted the fate of orally inhaled budesonide (highly soluble, mildly lipophilic) and fluticasone propionate (practically insoluble, highly lipophilic) in healthy subjects for: i) systemic and site-specific lung retention profiles, ii) aerodynamic particle size-dependent deposition profiles, and iii) identified the most impactful drug-specific, formulation-specific, and system-specific property factors that impact the fate of both the pulmonary and systemic concentration of the drugs. In summary, the presented multiscale computational model can guide the design of orally inhaled drug products to target specific lung areas, identify the effects of product differences on lung and systemic pharmacokinetics, and be used to better understand bioequivalence of generic orally inhaled drug products.

**Author summary:** Despite widespread use of available orally inhaled drug products (OIDPs), much is unknown regarding their optimal lung deposition, targeted delivery to specific lung regions, and the effects of various device, formulation, and physiological factors on deposition, absorption, transport, and clearance. In this study, we have presented a multiscale computational framework that integrates a full-scale 24 generation 3D lung model with distinct barrier regions spanning trachea, tracheobronchial, alveolar, and the terminal alveolar sacs with multiple other modules to track the OIDP levels (concentration) in both blood and pulmonary tissue regions. Along with validating the framework on two different inhaled drug types, we have also presented a sensitivity analysis to highlight the most impactful drug and formulation parameters, and therefore, potential optimization parameters to modulate lung selectivity and to better understand the pulmonary retention of drugs in distinct lung regions.

## Introduction

Respiratory diseases, such as asthma and chronic obstructive pulmonary disease, are among the leading causes of morbidity and mortality worldwide with an increasing burden on the healthcare and economies of all nations.[1] In most cases, the inhaled route of administration is the preferred method for delivering therapeutics for respiratory diseases, where treatment efficacy depends primarily on the quantity of drug deposited and distributed within the lung or at the specific site of action, which may be the upper or lower lungs.[2, 3] Compared to oral or intravenous routes, the inhalation route offers several advantages, such as: i) promoting high local drug concentrations directly at the site of action in diseased lung tissue, which may not be achievable efficiently by other routes,[4] ii) avoiding ‘‘first pass metabolism’’ of the liver which can greatly reduce drug concentrations before the drug reaches the systemic circulation,[5, 6] iii) rapid absorption (within minutes) due to large surface area of the lungs and high vasculature,[7] iv) favorable lung-selectivity (pulmonary efficacy/systemic safety ratio) that minimizes toxicity,[8, 9] and v) as an alternate route of administration for drugs that are effective systemically, but are not suitable for oral or intravenous administration, primarily due to low bioavailability.[10] At present, a large number of different types of inhalation devices exist in the U.S. market to deliver range of active pharmaceutical ingredients for the treatment of respiratory diseases.[11, 12] However, despite the widespread use of these orally inhaled drug products (OIDPs), much is unknown with respect to achieving optimal lung deposition, targeted delivery to specific lung sites, and the effects of various device, formulation, and physiological factors on deposition, absorption, transport, and clearance of these products. These limitations, along with the impracticality of obtaining human lung tissue concentration data of the delivered drug also make it difficult to evaluate and establish bioequivalence of potential generic products without a comparative clinical endpoint or pharmacodynamic bioequivalence study in the indicated patient population.[13]

Considering these inherent difficulties, *in silico* modeling offers a relatively efficient and cost-effective means of accelerating OIDP development. At present, many such *in silico* tools exist ranging from simple compartmental models to more complex physiologically based pharmacokinetics (PBPK) models. For example, Weber and Hochhaus provided a compartmental model for simulating human systemic pharmacokinetics of inhaled corticosteroids (ICSs) by incorporating selected physiological and formulation-related parameters,[14] whereas Boger et al.[15] and Hendrickx et al.[16] used semi-physiological compartment models to capture key features of both systemic and lung tissue pharmacokinetics profiles of multiple soluble bronchodilator drugs to rats and dogs and translated that model to predict the human plasma profiles. A different approach was employed by Gaz et al.[17] as an alternative to classic compartmental representations in which lung was further resolved to incorporate bronchial tree mucosa and smooth muscles to simulate hypothetical bronchodilator response in asthmatic conditions. An integrated approach of compartmental and PBPK modules has also been employed by Caniga et al.[18] to simulate rodent pharmacokinetics and its translation to humans. A similar, but more advanced, integrated model was recently employed by Hartung and Borghardt, that used a computational framework based on physiologically-structured population equations to integrate all relevant pulmonary processes mechanistically (deposition, clearance, dissolution, etc.), and evaluated against data from different clinical studies.[19] Commercially, the two main available PBPK software packages to model inhaled drug pharmacokinetics are Gastroplus™ (Simulations Plus Inc., Lancaster, CA, USA) which has mechanistic modules for regional deposition, dissolution, and permeation of inhaled drugs,[20] and SimCyp Simulator™ (Certara, Sheffield, United Kingdom) that has pulmonary delivery modules by reducing dissolution and epithelial permeation into a single first order process through a single pulmonary compartment. [For further information on available modeling approaches and their role in the development of OIDPs and devices, please refer to the reviews by Borghardt et al.,[21] Backman et al.,[22] and Walenga et al.[13]] Nonetheless, although these previous modeling efforts and available tools have proven to be useful, assessment of lung-selectivity has so far proven to be elusive and questions remain.

First, the predicted outcome of the drug in the systemic circulation is the result of pulmonary absorption (lung-to-blood) as well as gut absorption (swallowed fraction-to-blood), and hence, unbound concentrations of the drug in plasma alone may not be assumed to accurately reflect the target site-specific concentration in the lung without other justification.[4] Since the drug concentration in plasma and lung tissue is the result of parallel absorption from both gut and lungs and recirculation from blood-to-lung, a clear circulatory system (both systemic and pulmonary) must be defined in models along with the gut absorption models, systemic clearance, and region-specific mucociliary clearance (MCC) in the upper lung which is swallowed to the gut.[23–25] Second, in the physiological lung models, the heterogeneous nature of the lung with distinct differences between the tracheobronchial (also called conduction or central regions), alveolar regions (also called respiratory or peripheral regions), and alveolus (i.e., terminal alveolar sacs) should be made. In few previous modeling studies,[18, 19] the first two regions have been included in the modeling, but so far to the authors’ knowledge, no one has reported the separation of terminal alveolar sacs as a separate region. This is important because alveolar sacs are anatomically and physiologically distinct from the alveolar region due to the presence of a very thin air-blood barrier and surfactant layer. In previous studies, terminal alveolar sacs have been lumped as part of the alveolar region. Third, pulmonary drug disposition depends on a wide range of processes, including, the inhalation flow profile, distinct airway geometry, and particle size distribution (PSD) - all of which combine to produce a heterogeneous deposition pattern throughout the lungs. Hence, these parameters should ideally be part of the modeling effort before calculating/modeling further downstream processes of dissolution, MCC, and transport.

To address the aforementioned challenges, we present here a multiscale computational framework (Fig 1) that involves: i) our recently published full-scale 24 generation (Gen) 3D lung model with distinct barrier regions spanning trachea (Gen 0) to tracheobronchial (Gen 1-15) to alveolar (Gen 16-23) and to the terminal alveolar sacs (Gen 24);[26] ii) our previously published and modified computational fluid dynamics (CFD) module, called quasi-3D (Q3D), to calculate inhalation flow profile and PSD-based drug deposition,[27, 28] iii) a first-principles-based and lung region-specific dissolution and absorption module, iv) a tracheobronchial-region specific MCC module, and v) a gut absorption module, all connected to whole-body PBPK. Our simulation outcomes were validated on two distinct ICSs: budesonide (conditions specific to the Novolizer® device, which under normal inspiratory flow rates shows similar deposition of budesonide in the lungs of healthy volunteers as the Turbuhaler®)[29]) and fluticasone propionate (conditions specific to the Diskus® device). Finally, we also present a sensitivity analysis to highlight the most impactful drug and formulation parameters, and therefore, potential optimization parameters to modulate lung selectivity and to better understand the pulmonary retention of drugs in distinct lung regions.

**Fig 1.**
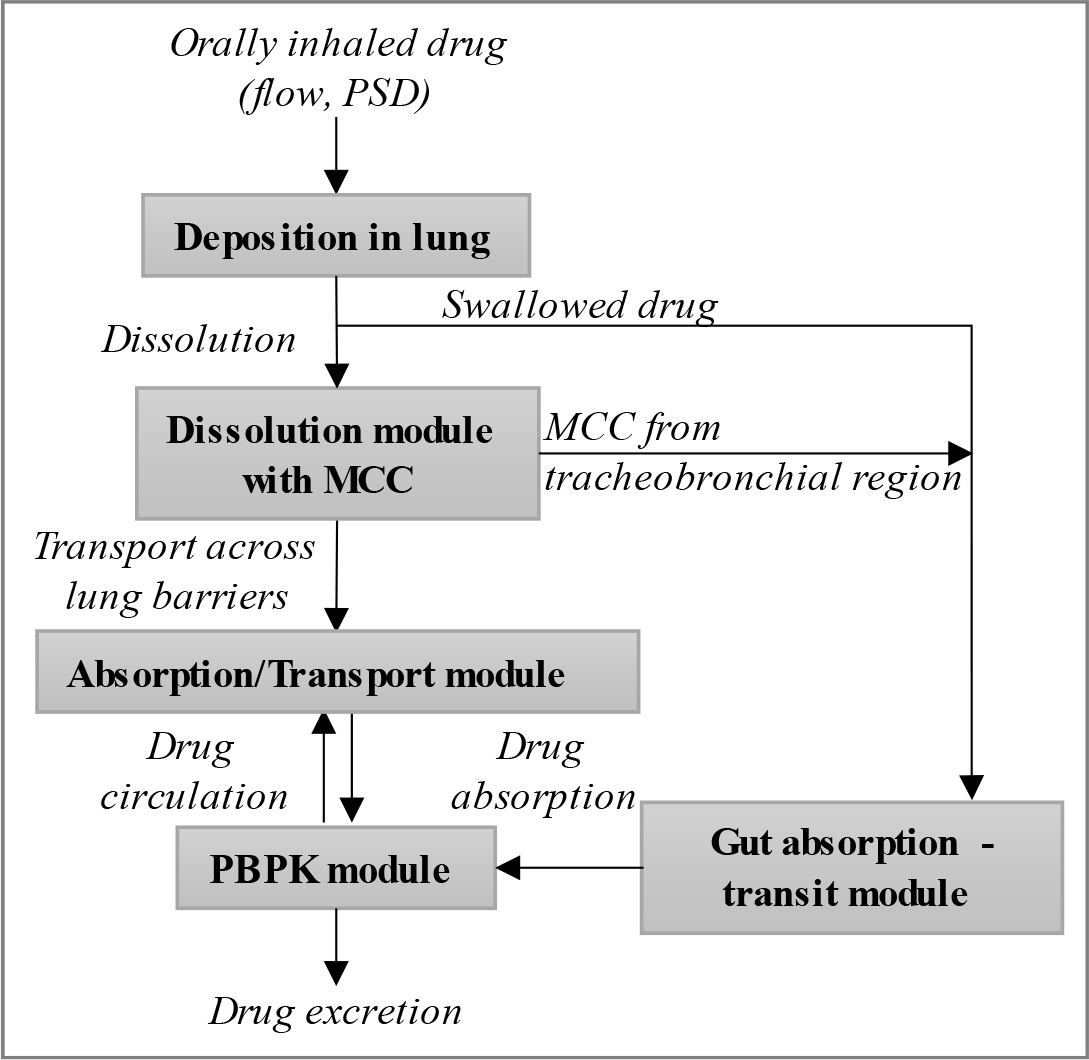
Computational framework to simulate orally inhaled drugs. Computational modules are shown in blocks whereas the pulmonary processes are in italics.

## Models

### Simulated drugs

The goal of this work is to develop and validate a mechanistic pulmonary pharmacokinetics model that can capture most of the relevant physiology and biophysics involved in inhaled drug pathway starting from breathing profile and drug PSD to final outcomes of drug concentration in systemic plasma and pulmonary tissue. For model validation, we have selected two different types of ICSs - budesonide and fluticasone propionate. In terms of physicochemical properties, budesonide has relatively high aqueous solubility and is mildly lipophilic, whereas fluticasone propionate is practically insoluble and is highly lipophilic.[23,30,31] These differences impact dissolution, absorption, luminal clearance, and lung retention time, which in turn influence the final, and distinct, systemic and pulmonary tissue profile of these drugs.

### The Quasi-3D (Q3D) lung model

To determine the deposition profile of these drugs, we employed CFDRCs *in house* developed Q3D technique using a full-scale 3D lung model.[27,28,32,33] The dimensions of the lung model correspond to the 50th percentile adult U.S. male (172 cm in height, 70 kg mass).

Q3D method. In many biomedical and engineering problems, physical processes occur in networks of pipes/tubes, cables, wires, or other one-dimensional (1D) structures. The best examples are the human vascular system, lymphatic network, neurons with a network of dendrites and axons, microfluidic channels in biochips, and of course the case of airflow transport in the lung airways. Full-fledged 3D computational simulations of such large tubing structures are possible for some cases such as inhaled particle transport and deposition in the upper lung airways (wherein the total physical time is in the order of seconds).[32, 34] However, 3D computational simulations that require a physical time scale greater than several tens of seconds (or more) are computationally demanding and depending on availability of high performance computational resources, may not feasible. This is particularly relevant in simulating the particle transport/deposition in the lung airways that require several breathing cycles. A 1D model of a tubing network distributed in a 3D space is well suited to solve such a problem, as previously shown by authors.[26–28] The major advantages of this approach are the ease of model setup, high computational speed, simple visualization of results, and an easy link to compact models such as spring/mass/damper devices, valves, pumps, controllers, and 0D compartmental models. This method is referred to as the Q3D model, since it solves for all the 3D flow variables of {u,v,w,p}x (unlike 1D models) while maintaining the fully developed wall boundary condition. Details on the Q3D creation, its accuracy, its speed, solution accuracy (including in the context of the flow in the human lung), problem setup, robustness, details on the flow solver, the assembly of the matrices, the spatial and temporal schemes, and the modeling of the turbulent stresses are available in Kannan et al.[27] Lung model. Most known lung models typically contain the geometry of only the first 6-9 airway branch generations due to the resolution of available imaging data that does not permit accurate visualization of smaller branches in further generations. A full 24 generation lung model of an adult male human was developed for this study. Unlike previous full lung models,[35, 36] the newly developed model was used to simultaneously simulate (i) flow transport simulations, i.e., inhalation and exhalation simulations and (ii) aerosol transport and deposition simulations, over several breathing cycles. In this section, we will briefly describe the process for: i) extending the Q3D lung which was extracted from the Zygote stereolithography (STL format) to the end of the tracheobronchial limit (i.e., Gen 0-15), and ii) constructing the “sac-trumpet” like control volumes at the end of the tracheobronchial exits to mimic the alveolar region (Gen 16-23) and terminal alveolar sacs (Gen 24).

As the first step, we extended the Zygote lung model to the end of the tracheobronchial limit. The lung lobes provide the outer boundary for the extension process. Fig 2A-B, shows the lung lobes, enclosing the original Q3D lung with and without the lobes (created from the Zygote lung model – details provided on the zygote website: https://www.zygote.com/poly-models/3d-male-systems/3d-male-respiratory-system).

**Fig 2.**
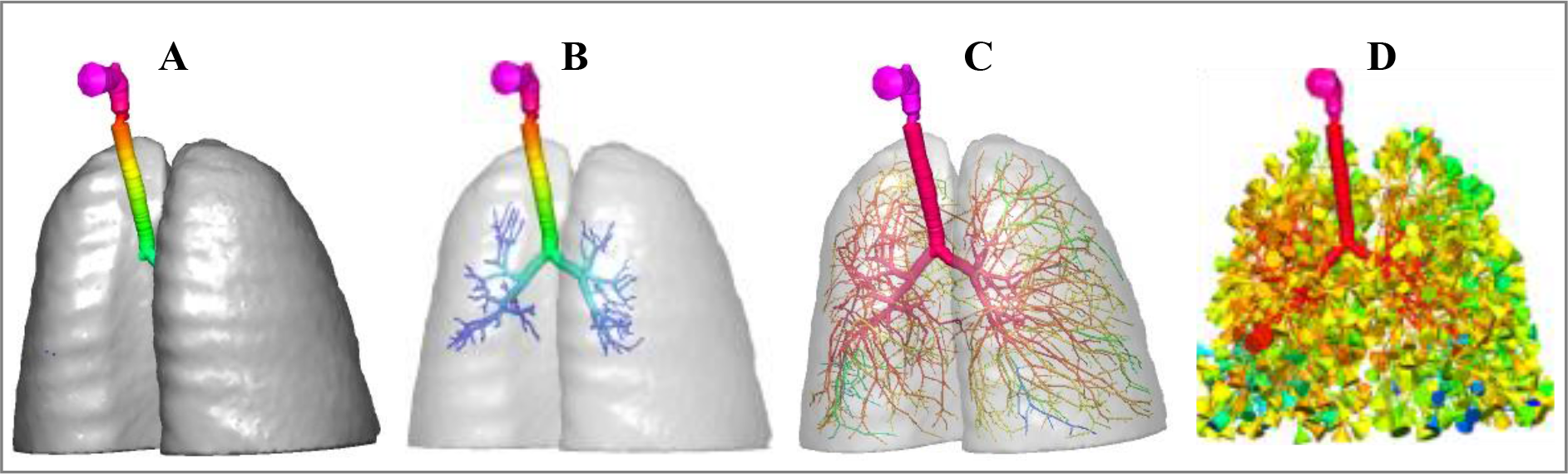
The development stages of the full 24 generation 3D lung. The stages show the original imaging-based human Zygote lung with tracheobronchial extensions limit up to generation 6-9 in opaque lobes (A) and (B) transparent lobes; The extension of tracheobronchial limit up to generation 15 (C); and the whole lung with extensions up to generation 23 and sac-trumpet representation of terminal alveolar sacs (generation 24) (D). The sac-trumpet representation of the whole lung is colored by higher to lower pressure (pink>red>yellow>green>blue) for an inhalation flowrate of 5 L/min.

Next, we adapted the algorithm of Karch et al.[37] to extend the current Q3D airways to the end of the tracheobronchial limit and implemented sac-trumpet like control volumes at each of the tracheobronchial outlets (Fig 2C). Fig 2D shows the complete Q3D lung, i.e., after the insertion of the sac-trumpet control volumes. The total functional residual capacity (FRC) in the tracheobronchial section (excluding the mouth, nasal, oral, laryngeal, and pharyngeal sections) is around 165 cc. This volume is similar to values presented in the literature, such as Pichelin et al.[38] provides a value of around 130 cc for a 1.81 m tall male human, whereas the Weibel model value is ∼155 cc.[39] The overall tracheobronchial lateral surface area of this generated lung is ∼1996 cm^2^. In general, it is difficult to recreate a lung model whose areas and volumes both match that of the real lung because the surface of the airways (and especially the terminal alveolar sacs) of the actual lung is non-smooth and folded to enhance the lateral surface area. The tracheobronchial lateral surface area for the real human lung is 2471 +/- 320 cm^2^ as per the experimental measurements of Mercer et al.[40] The FRC of the developed whole Q3D lung model is 2611 cc.

### Lung barriers

The above developed lung model is then modified to include various generation-specific barrier layers. Overall, the airway barrier model simulates the MCC (axial direction, from tracheobronchial→throat→gut) and trans-mucosal transport (radial direction, from airway lumen→lung tissue→blood), as well as the dissolution of deposited drug on the airway walls. As described in the Introduction section, the existing models of pulmonary barrier models use a compartmental approach, in which the pulmonary wall is divided into two “axial” segments: tracheobronchial and alveolar. Each segment consists of several layers (starting from the lumen): mucosal gel and sol (together called mucosa), epithelial layer, stroma layer with embedded airway smooth muscle cells and immune cells, and the pulmonary endothelial layer. It is important to note that due to the heterogeneous nature of the human lung, the barrier dimensions for these layers change from generation to generation (Fig 3). For example, the heights of epithelial cells range from 50-80 µm in the trachea[10, 41] and gradually taper down to less than 0.5 µm in the alveolar sacs.[42] Since it is not possible to obtain experimental values of the changes in these barrier dimensions for all the individual 24 generations, few previous studies have ”lumped” them together in tracheobronchial and alveolar regions with approximate average dimension values. [Authors suggest the review articles by Frohlich et al.[41] and Patton and Byron[10] for more discussion in human lung barrier dimensions.]

**Fig 3.**
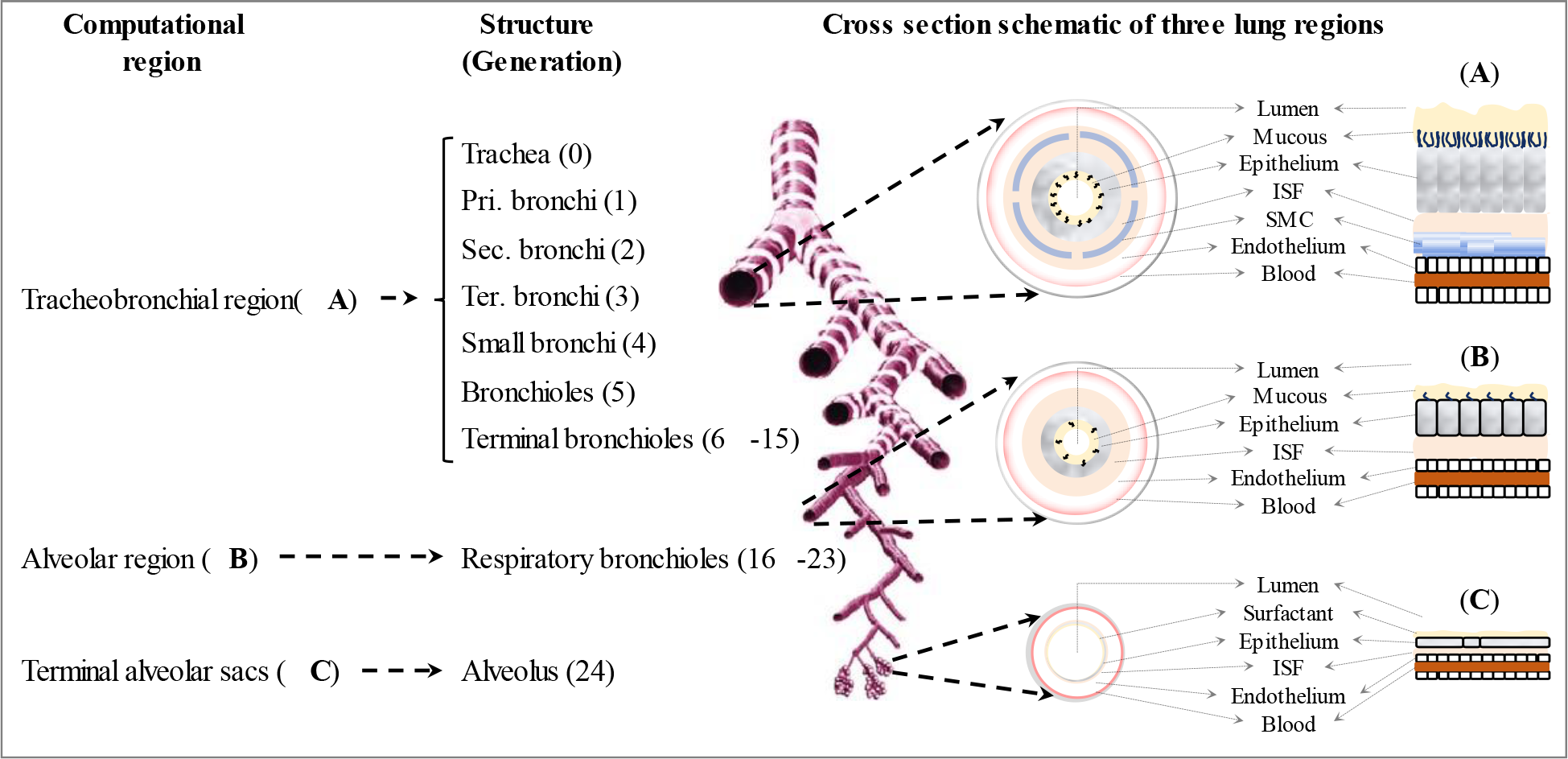
Schematic of the three different lung regions and barrier layers modeled in this study. Dimensions are not to scale. [SMC = smooth muscle cells; ISF = interstitial fluid].

The optimized values of these dimensions in the two lumped compartments and the subsequent ordinary differential equations to describe the transport of drugs through these barriers were first formulated by Yu and Rosania.[43] Briefly, this model lumps the first 16 lung branches (Gen 0-15) into the conducting region (i.e., tracheobronchial) and the last 9 generations (Gen 16-24) into the respiratory region (i.e., alveolar). Such lumped models are based on the approximate structural and functional differences between the conducting and respiratory regions. In absence of generation-specific experimental data, this also greatly simplifies the model for drug/particle transport and facilitates a fast and easy simulation of their transport across the air (lumen)-to-blood barrier.

In this work, we have adapted the above-described lung barrier model of Yu and Rosania into the Q3D framework to simulate the dissolution and transport of the drug across the airway barrier in the entire airway tree. In this model, at each airway axis position the air-to-blood barrier, starting from the mucosa to the plasma in lung tissue, is radially divided into several layers representing each type of cells in the tissue. A set of ordinary differential equations is solved in each layer to simulate drug dissolution, diffusion, convection (in the mucosal layers), binding and absorption into pulmonary circulations. Some subcellular organelles such as lysosomes and mitochondria are also modeled as sub-compartments in each layer for their role in determining drug pharmacokinetics. Also, since the alveolar sacs have significantly different properties, due to the very thin barrier and presence of surfactant or surface lining liquid – SLL,[8] from the general alveolar region, in our model we have designated the alveolus as a separate compartment, called the terminal alveolar sacs region. Though physiologically, the terminal alveolar sacs can start as early as generation 18, the majority of total alveolar sac volume comes from generation 24,[38] hence for simplicity the presented model only considers the terminal alveolar sacs as part of generation 24. The overall schematic of our barrier model of the different layers and their lung region-specific description is provided in Fig 3. The other main parametric changes include: i) modified permeability of terminal alveolar sacs region to add surfactant effects, ii) modified permeability of the alveolar region, iii) modified dissolution coefficient in the mucosa (which is present in Gen 0-23) and surfactant, and iv) recalibrated barrier thicknesses in each of the three defined lung regions.

The model also accounts for key physicochemical properties of the transported molecules that are required as model inputs, including: i) logP, ii) blood-to-plasma ratio (B2P), iii) free fraction of the drug in plasma (fu), iv) particle density, v) diffusivity and solubility in a water-like fluid (mucosa), vi) tissue barrier permeability, vii) the deposition distribution, viii) the drug valency, ix) the organ clearance rates for lung, liver, and kidney, and x) partition coefficients.

These parameters determine the transfer rates, i.e., the rate at which the solid drug is converted to the molecular form and then absorbed into the plasma/tissue.

The modified barrier thicknesses (biological parameters) and the key physicochemical parameters (drug parameters) were finalized using rigorous optimization of the model’s systemic output (plasma concentration of inhaled drugs) and its match with the known experimental data. Whenever possible, the base range of these parameters were within the known bounds of experimental values. For example, the SLL thickness in the terminal alveolar sacs has been reported with values of 0.01–0.08 μm by Olsson et al.,[44] 0.1–0.2 μm by Wauthoz and Amighi,[45] 0.07 μm by Patton and Byron[10] and 0.3 μm by the National Research Council.[46] Hence, to optimize the value of SLL we used the lowest (0.01 μm) and highest (0.3 μm) reported range for this parameter and iteratively optimized it while keeping other parameters constant and picked the final value that gave us the most optimum simulated budesonide and fluticasone propionate pharmacokinetics area under the curve (AUC) compared to experimentally known budesonide and fluticasone propionate AUC in healthy human subjects.

Other biological parameters were similarly optimized by collecting the reported min-max range in other studies.

The previously published values and our final optimized values of the parameters are shown in Table 1.

**Table 1.**
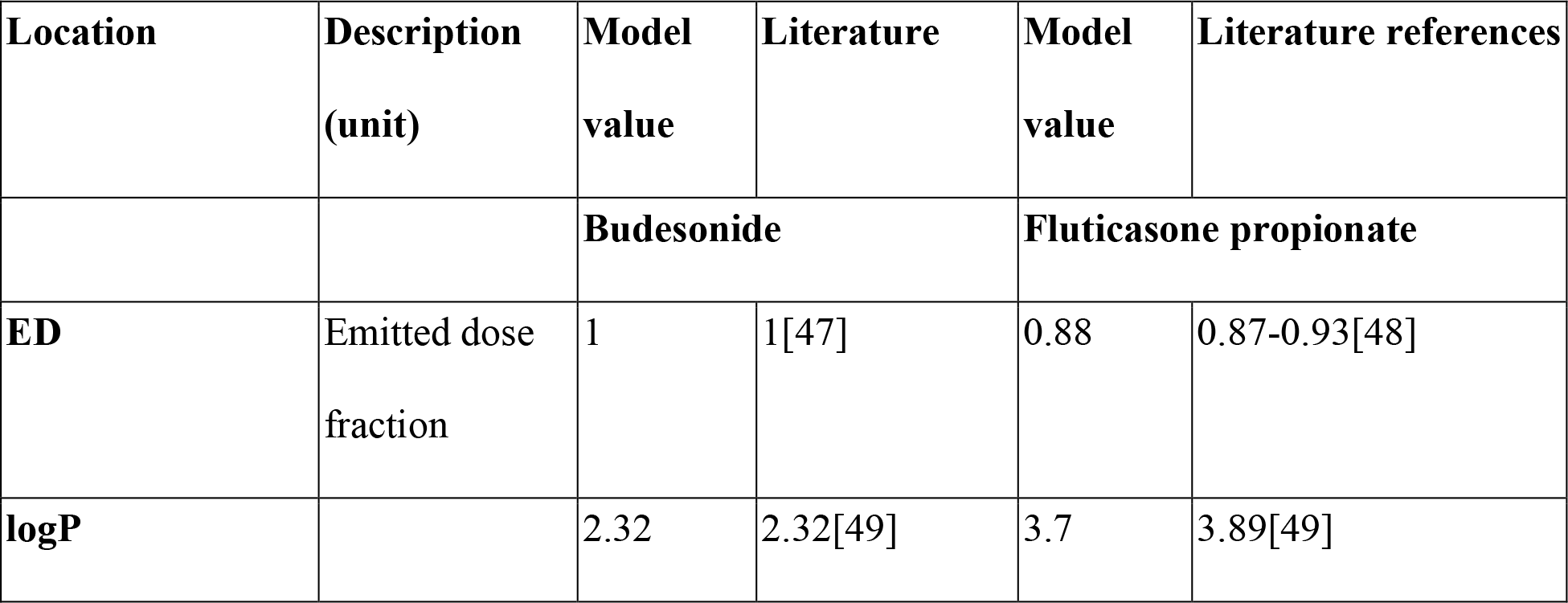
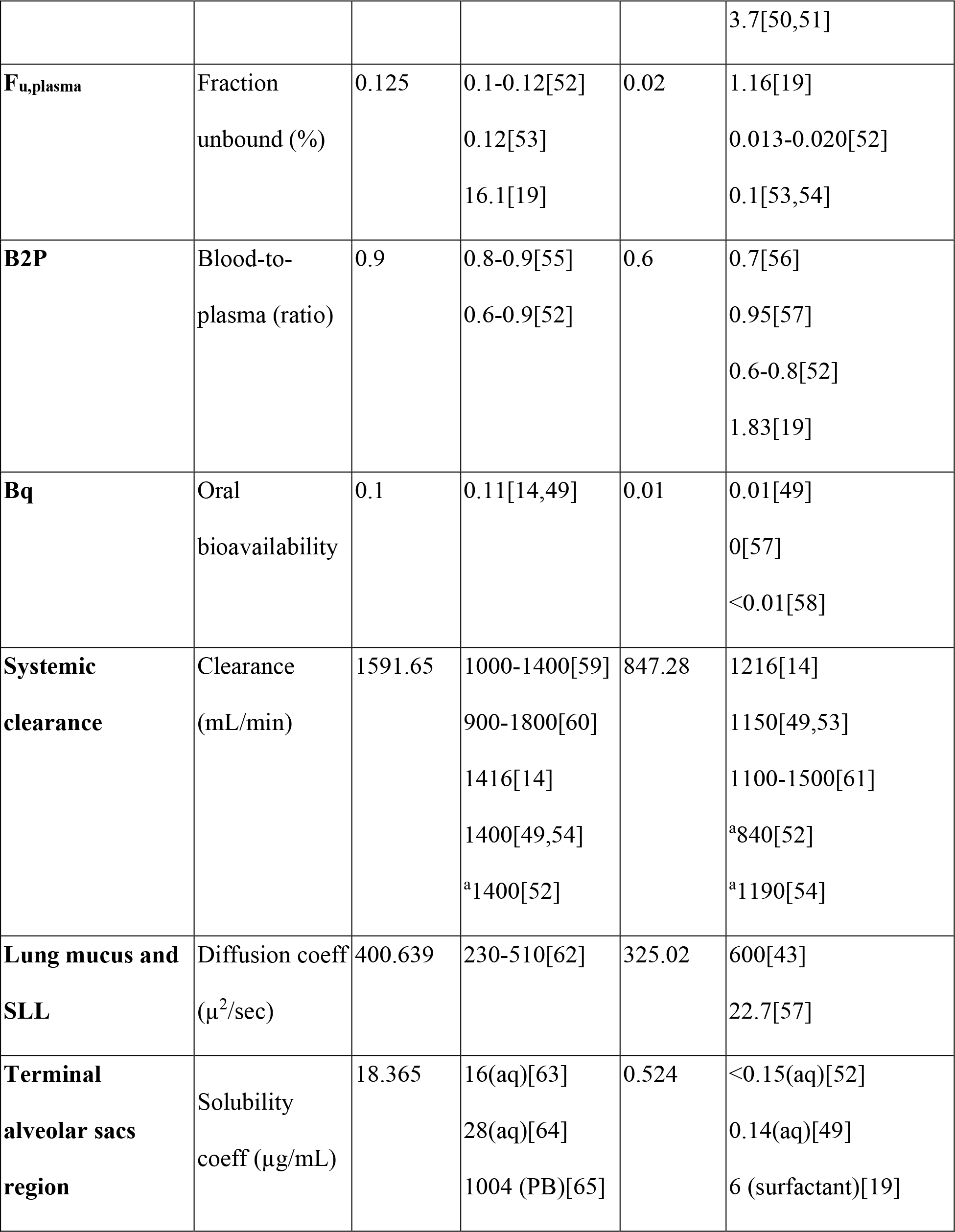
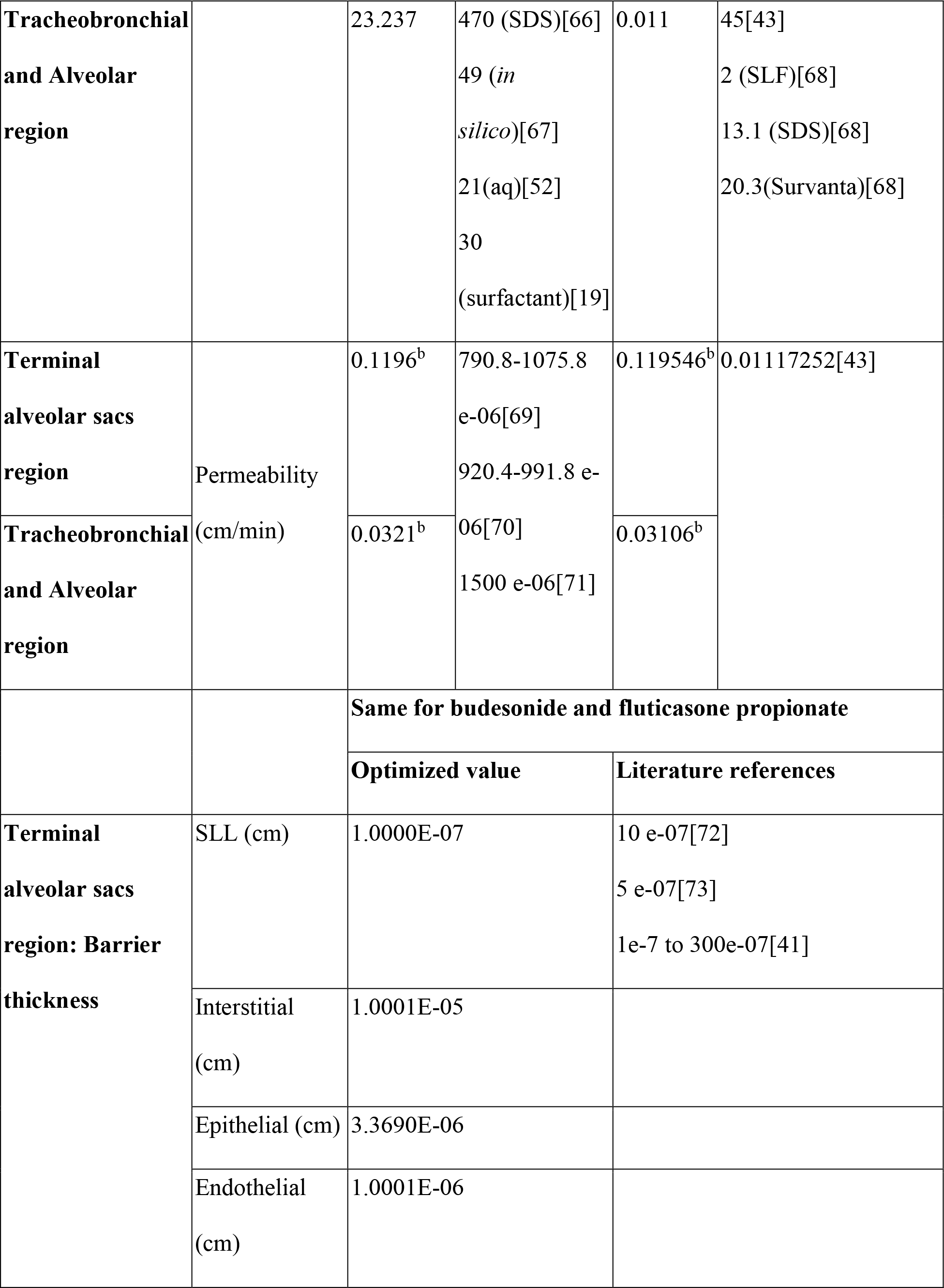
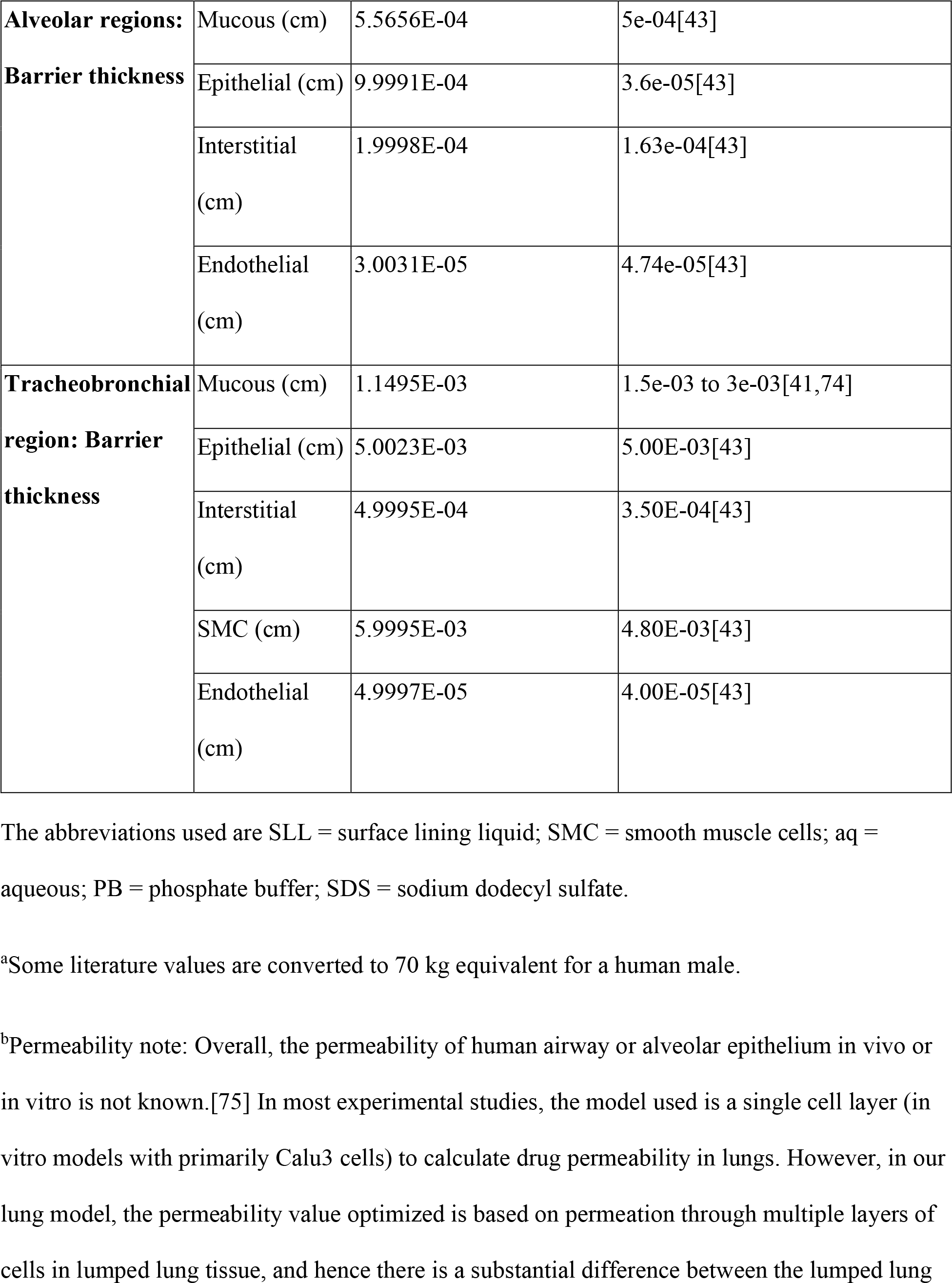
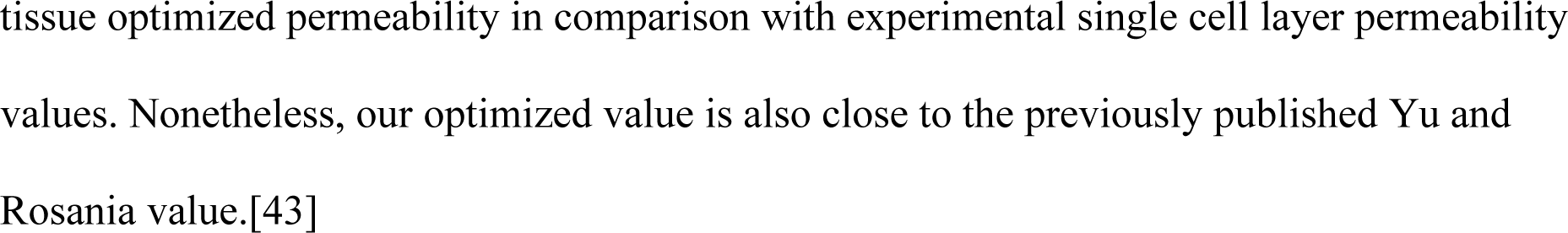
Drug specific parameters and biological parameters (lung barrier thickness) used in the model for drugs budesonide and fluticasone propionate.

### Equations for barrier transport

Only neutral and ionized drug in the aqueous phase is allowed to transport across radial airway barrier layers. The neutral drug transport is passive and driven by the activity difference in two neighboring compartments and follows Fick’s first law [Activity here is collective effect of the terms contributing to the total mass flux]. The ionized drug transport is driven by the electrochemical potential difference and described by the Nernst-Plank equation. The list of all the transport flux across all the barriers in individual lung regions is rather large and the authors recommend the Yu and Rosania study for further details.[43] Here, we are demonstrating the key idea of modeling transport flux and of being linked to other modules in the computational platform in the following way: Consider the drug flux between the endothelial compartment (compartment 7) and the plasma compartment (compartment 8) in the tracheobronchial region (i.e., airway region (AW) as per Yu and Rosania’s convention), i.e., the transport of a neutral drug from endothelial barrier to systemic blood at a specific lung generation. Since the neutral drug transport is passive and driven by the difference of neutral drug activity in the aqueous phase in two neighboring compartments it follows Fick’s first law:

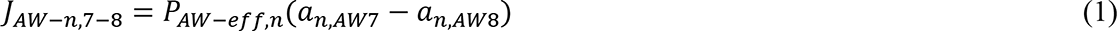

Here, JAW-n,7-8 is the transport flux on a unit surface area of the neutral drug across the barrier between (compartment) 7 and 8 in the AW region; it has a unit of drug mass over time over an area such as µg/min/cm^2^; an,AW7/8 is the neutral drug activity in the aqueous phase in compartment 7 or 8 in the airway region; and PAW-eff,n is the drug permeability of the barrier. The total mass flux rate across the barrier JAAW-n,7-8 (with a unit of drug mass over time such as µg/min) is obtained by multiplying the total surface area for drug transport:

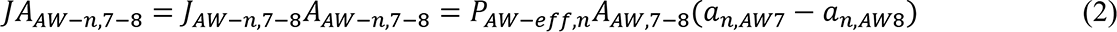

where AAW7-8 is the area of the barrier.

In contrast, the ionized drug transport is driven by the electrochemical potential difference and described by the Nernst-Plank equation. Hence, the net flux on a unit surface area of ionized form drug is described by the following equation:

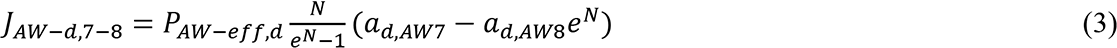

Here, JAW-d,7-8 is the transport flux of the ionized drug across the barrier between compartment 7 and 8 in the AW region; it has a unit of drug mass over time over an area such as µg/min/cm^2^. ad,AW7/8 is the ionized drug activity in the aqueous phase in compartment 7 or 8:

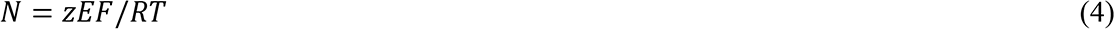

where, z is the electronic charge of the ionized drug molecule, E is the membrane potential, F is the Faraday constant, R is the universal gas constant, and T is the absolute temperature.

The total mass flux rate of the ionized form drug across the barrier JAAW-d,7-8 (with a unit of drug mass over time such as µg/min) is obtained by multiplying the total area for drug transport:

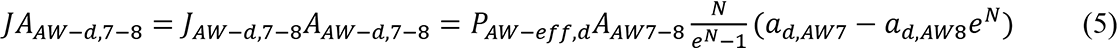

The overall net flux on a unit surface area JAW,7-8 is the sum of the neutral drug flux and the ionized drug flux. The fluxes between the other compartments are constructed similarly. These base equations were used in our modeling approach to describe the drug flux between three regions: tracheobronchial, alveolar and terminal alveolar sacs.

### Pulmonary drug deposition

For budesonide deposition studies in the above-described lung model, we used Novolizer® dry powder inhaler (DPI) device-specific conditions.

Previously, we have used and published an Euler Lagrangian (E-L) methodology to simulate the budesonide deposition for the same device using CoBi tools.[32] We used seven bins for the particle sizes, an aerosol velocity of 30 m/s, and a spread half-angle of 10.5o. More details related to the simulation setup, including the particle diameters, the distribution of the particles in the binds, the flow conditions, the spread angle, etc., can be obtained from that study. In the present work, we have used a Euler Euler (E-E) formulation in the Q3D framework. This is expected to be much faster than the E-L simulations due to: i) use of larger timesteps for the aerosol species, as opposed to smaller timesteps for the particles, ii) number of degrees of freedom being much smaller in the Q3D, as opposed to the large CFD mesh, and iii) faster solver convergence in the Q3D model, compared to the CFD models, due to the absence of skewed and high-aspect ratio cells.[32, 34] Hence, the current methodology can be used for simulating longer physiological responses, such as forced exhalation and secondary (multiple) breathing cycles.

For fluticasone propionate deposition, we used Diskus DPI device-specific conditions.

For both drugs, we used a starting dose (mass) of 1 mg inhaled drug and used a standard breathing profile (tidal volume = 0.5 liters, inhalation time = 3 seconds, and exhalation time = 3 seconds), after the initial forced inhalation. The comparison of discretized PSD and flow profile used in these device-specific simulations is shown in Fig 4 and are obtained from the experimental studies.[47,76,77]

**Fig 4.**
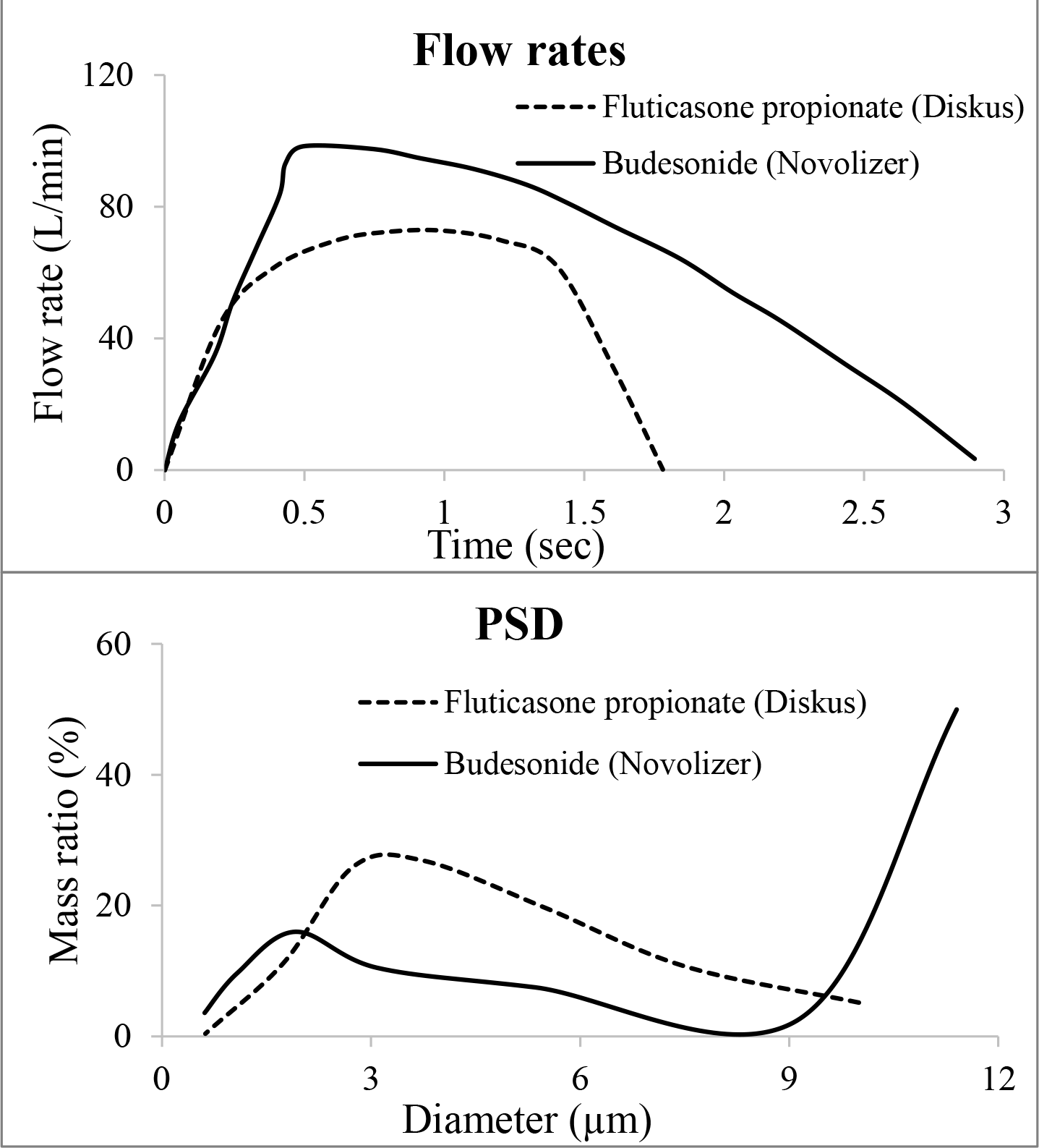
Flow rates and particle size distribution (PSD). The inhalation flow rate (top) and PSD (bottom) used in the simulations of budesonide and fluticasone propionate.

The aerosol transport equations, probabilities of deposition and the mesh independence studies are provided in the Supplementary Information (SI) document.

### Mucociliary drug clearance

Due to the continuous beating of cilia in the upper lung (especially the tracheobronchial region), the mucus layer covering the cilia moves drug microparticles towards the pharynx in a coordinated manner to effectively clear the deposited particles out of the airways, called MCC.[78] Therefore, it is necessary to account for both the dissolution and MCC processes to accurately characterize the particle deposition dynamics and patterns in the airways.

The MCC convection flow rate in the conducting airway is obtained from the literature.[79] The mucus velocity mV has the following fitted formula:

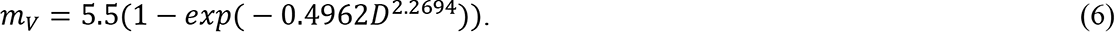

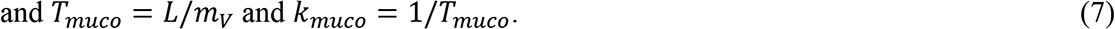

*D* is the airway diameter, *Tmuco* is the residence time, *L* is the airway length, *kmuco* is the rate constant of the MCC due to cilia beating.

### Pulmonary drug dissolution

The Noyes-Whitney equation is used to describe the dissolution process.[80, 81] In the computational platform, we use the following equations for drug dissolution in every control volume, assuming spherical geometry of dry particles of the drug:

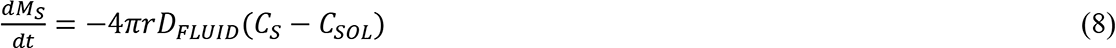

Here MS is the undissolved drug mass in the compartment, CS is the drug solubility coefficient in the compartment, CSOL is the local dissolved drug concentration and r is the microparticle radius. The above equation is applied in each generation of the lung model.

However, the solubility coefficient values (Table 1) for both drugs are different in the terminal alveolar sacs due to the presence of surfactant in the SLL.[68]

### Whole-body drug distribution and clearance

The drug reaching the outer barrier layer crosses the pulmonary epithelium and is absorbed into the systemic circulation, which is simulated as a whole body multi-compartmental PBPK. The organs are represented as well-stirred reactor 0D compartments. [Details of whole-body PBPK framework can be obtained from authors earlier publications].[28, 82] The drug concentration equation in perfusion rate-limited organs including fat, brain, bone, heart, muscle, skin, thymus, stomach, pancreas, spleen, and other is given by:

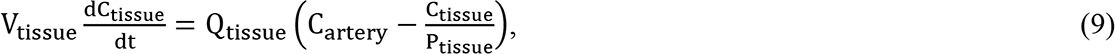

where Vtissue is the tissue volume, Ctissue is the drug concentration in tissue, Qtissue the perfusion rate, Cartery is drug concentration in the arterial blood, and Ptissue is the tissue distribution coefficient.

The drug concentration equation in the permeability rate-limited organs, the liver in particular, is given by:

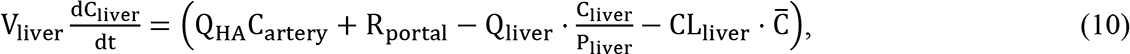

where QHA is the flow rate in the liver artery, R_portal_ is the drug entry rate into the liver via the portal vein which collects the blood from the stomach, pancreas, spleen, small and large intestine, Qliver is the flow rate in the liver vein and is the sum of the flow rates in the liver artery and the portal vein, ^C̅^is the average drug concentration from the liver artery and the portal vein and CLliver is the liver clearance rate. R_portal_is given by:

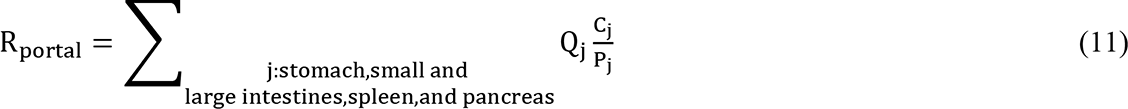

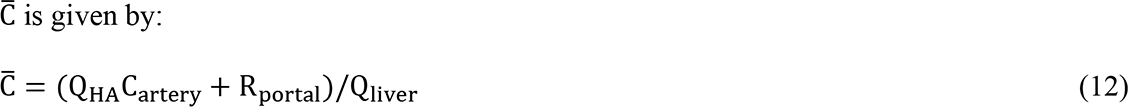

The drug concentration equation in the kidneys is given by:

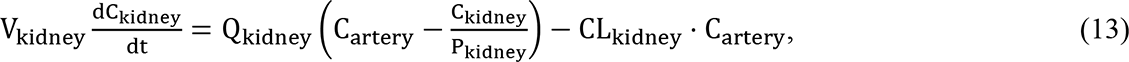

where CL_kidney_ is the kidney clearance rate.

The drug concentration in the venous compartment is given by:

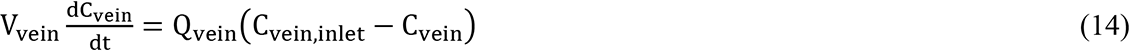

where C_vein,inlet_ is the average drug concentration in blood entering the vein from tissues.

The drug concentration in the artery is given by:

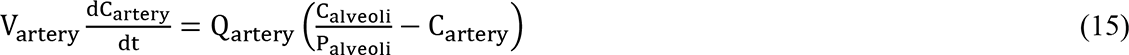

Here Qvein and Qartery are equal to the cardiac output.

## Results

### Drug deposition

Post-inhalation, fractions of drug particles are deposited in the various regions of the respiratory system following contact with the lung mucous/SLL. This process is influenced by several factors related to the particles’ physicochemical properties as well as physiological and anatomical features of the lungs.[83] The main physical processes determining respiratory drug deposition are impaction, sedimentation, and diffusion, which in turn are influenced by particle size, shape and density, as well as breathing patterns, and lung anatomical and physiological parameters. Following this general pattern, Fig 5 shows the steady-state deposited mass (in µg), for three selected diameter test-cases in our model: large (11.4 µm), medium (3.08 µm), and small (0.613 µm). The inhaled mass is normalized to 1 µg (in each diameter bin). The main, and expected (more deposition of smaller particles in the deeper lung, and vice-versa), observations are: i) larger particles (11.4 µm) get primarily deposited in the mouth-throat and glottis regions and we observed very little deposition in the lower lung (alveolar region) and the alveolar sacs for these particles, ii) we observe some inertial deposition for the medium sized (3.08 µm) particles that primarily get deposited in the upper lung regions and a small fraction in the alveolar sacs region, and iii) for smaller particles (submicron), we observe significant deposition in the terminal alveolar sac region. Overall, the deposition percentage values (ratio of the mass deposited in that region to the dosage mass) for both the tested drugs in the different lung regions are provided in Table 2. This is in direct correlation with the device-specific PSD data that was used as input in the model, for example, budesonide (in comparison with fluticasone propionate) has more particles in the submicron range and also in the particles that are larger than 10 µm, and hence, shows slightly larger values of deposited fraction in terminal alveolar sacs for these submicron particles, and in the mouth-throat and tracheobronchial regions for the larger particles, as compared to fluticasone propionate. In contrast, fluticasone propionate has more drug particles in the range between 3-9 µm that can bypass the upper lung generations but cannot travel all the way to the terminal alveolar sacs, and hence, has a higher predicted value of deposition of drug particles in the alveolar region.

**Fig 5.**
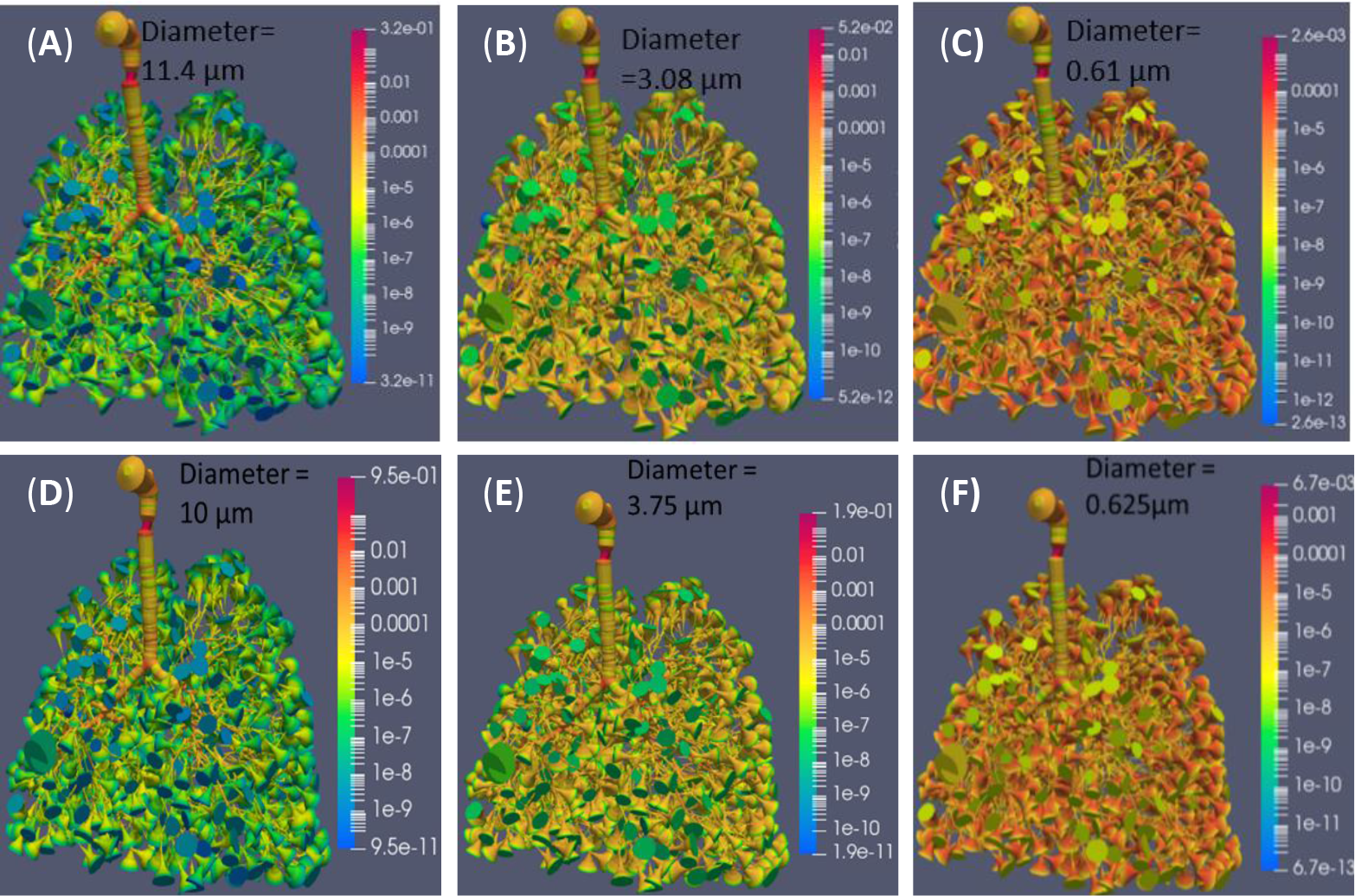
Deposition pattern. The steady-state deposited mass (in µg) is shown for three selected particle sizes to highlight the size-based inhaled drug deposition for budesonide (A-C) and fluticasone propionate (D-F). Inhaled mass is normalized to 1 µg. Red to blue shows higher to lower deposition.

**Table 2.** The deposited mass (% of metered dose) in different lung regions for budesonide (Novolizer) and fluticasone propionate (Diskus).

| **Lung Region** | **Lung Generation** | **Budesonide (%)** | **Fluticasone propionate (%)** |
| --- | --- | --- | --- |
| **Mouth-piece** |  | 11 | 12 |
| **Mouth-throat** |  | 40.2 | 33.98 |
| **Tracheobronchial** | Gen 0-16 | 29.46 | 17.16 |
| **Alveolar** | Gen 16-23 | 15.43 | 33.87 |
| **Terminal alveolar sacs** | Gen 24 | 3.91 | 2.99 |

Using this Q3D E-E method, the total lung deposition fraction (without trachea, which was 1.8% of the metered dose) for budesonide was predicted to be 47% of the metered dose. This falls outside of the experimental mean value of 36.5% (recalculated from the 9.4 – 41% (median 32.1%) as described by in vivo γ scintigraphy studies of budesonide deposition by Newman et al.[29] The data from the highest peak inspiratory flow rate (PIFR) of 99 LPM were used for comparison as this flow rate value is most consistent with the intended operating flow rate of the simulated device at a standard 4 kPa pressure drop. The study by Newman et al. has further provided the deposition fractions of central, intermediate, and peripheral lung regions; however, we have refrained from making simulation comparison with these values due to lack of consistence in regional split of lung between the two studies. For example, in computational models, the data are usually analyzed in terms of fractional deposition in tracheobronchial and alveolar regions that are designated a specific generational numbers, while the physiological lung regions are mixtures of generations as recently highlighted by Olsson et al.[84] Moreover, there are additional differences between the presented modeling protocol and the γ scintigraphy experiments that could account for the total lung deposition difference. This includes the use of male lung scan-based model in simulations (the γ scintigraphy study subjects comprised of both male and female test subjects) and the input PSD profile in modeling (the PSD profile of the DPI device used in γ scintigraphy study is not provided). Although the PSD profile was not provided in the Newman et al. [29] study, the fine particle fraction (FPF) was given, which can affect the regional deposition. The FPF provided by Newman et al. [29] is 34.9% +/- 5.1%, whereas the FPF ranges from 40-47.5% in the present case (considering the cut-off for the FPF as 5 µm). The difference in FPF between the simulations and γ scintigraphy experiments may explain why the prediction for total lung deposition was higher than the in vivo data.

### Systemic drug concentration

As the first step, before predicting the systemic drug concentration, we identified and analyzed the appropriate clinical systemic pharmacokinetics datasets for both drugs. The five available datasets for each of these drugs are shown in Fig 6. It is important to note that: i) all the selected experimental datasets are based on healthy human subjects, as our developed lung model is based on healthy lungs, ii) for comparison, all the datasets are normalized to 1 mg dose, iii) all datasets used only single drug types to avoid synergistic/antagonistic effects, iv) due to the variation in experimental datasets (for example, a slight difference of 1 mg and 1.2 mg of inhaled budesonide dose create a dose-normalized difference of two-fold between maximum plasma concentration (Cmax) and AUC from time zero to infinity (AUC0-∞) values while maintaining the overall shape of the pharmacokinetics plots),[85, 86] our goal is primarily to compare with the *average* time-concentration profile of the collected experimental datasets, and iv) the comparison matrices of model versus experiments were evaluated in terms of the visual relative shape of the pharmacokinetic plots, as well as, the quantitative pharmacokinetic parameters (Cmax, time to Cmax (Tmax), and AUC0-8hr).

**Fig 6.**
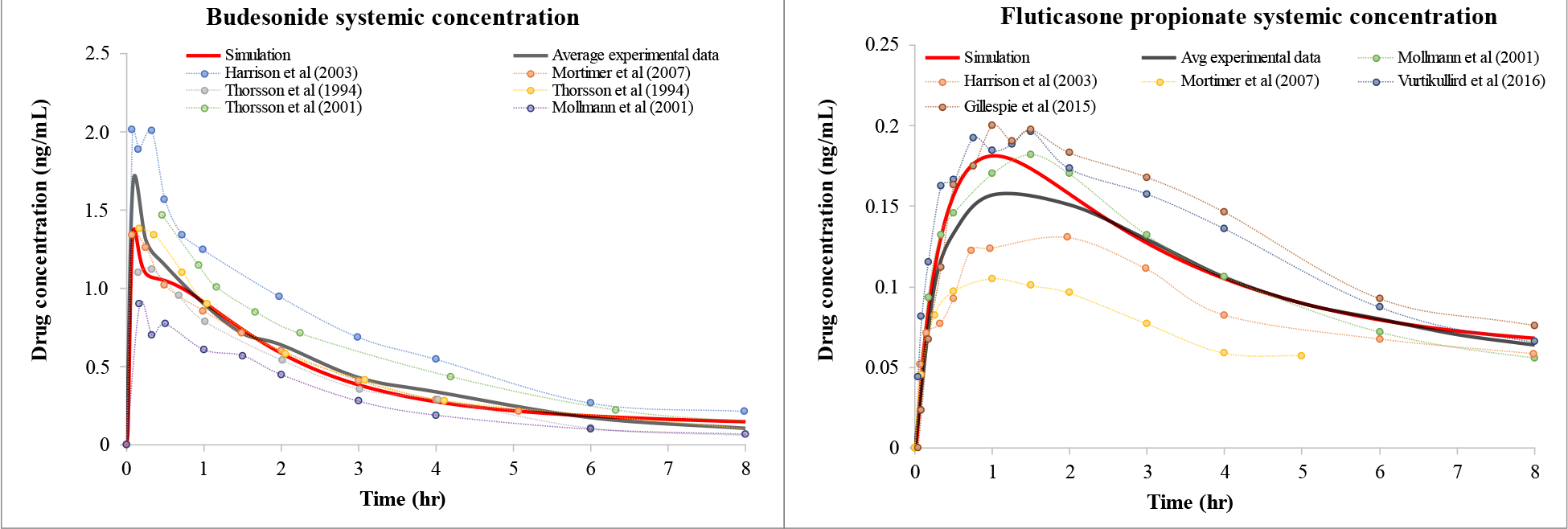
Plasma (systemic) concentration-time profiles. The simulated concentration-time profiles are shown for administration of 1 mg of budesonide inhaled with Novolizer and fluticasone propionate inhaled with the Diskus devices. Clinical data points: digitalized raw data from multiple references normalized to 1 mg.[85–89] The black line shows the average of all clinical data points and red line is the simulation predictions. [Note: Two data points from Thorsson et al. (1994)[88] are from two different datasets in the same article].

Fig 7 shows the predicted plasma systemic pharmacokinetics profile of budesonide and fluticasone propionate in comparison with the clinical experimental data of healthy patients. For both cases, the dose-normalized data from literature were in agreement, where the simulation pharmacokinetic outcomes closely matched the average experimental data in terms of AUC0-8hr values. Cmax of budesonide is slightly underpredicted, whereas Tmax was underpredicted in both cases, but all values were well within the experimental range as shown in Fig 7 and Table 3-4. Noticeably, our model was able to predict the bi-phasic (a peak and a bump) budesonide response, which was shown in some experimental data to occur within 20 minutes of drug inhalation. This has previously been observed in the in vivo pharmacokinetics studies of Mollman et al.[86] and Harrison et al.[85] and is further described in the Discussion section.

**Fig 7.**
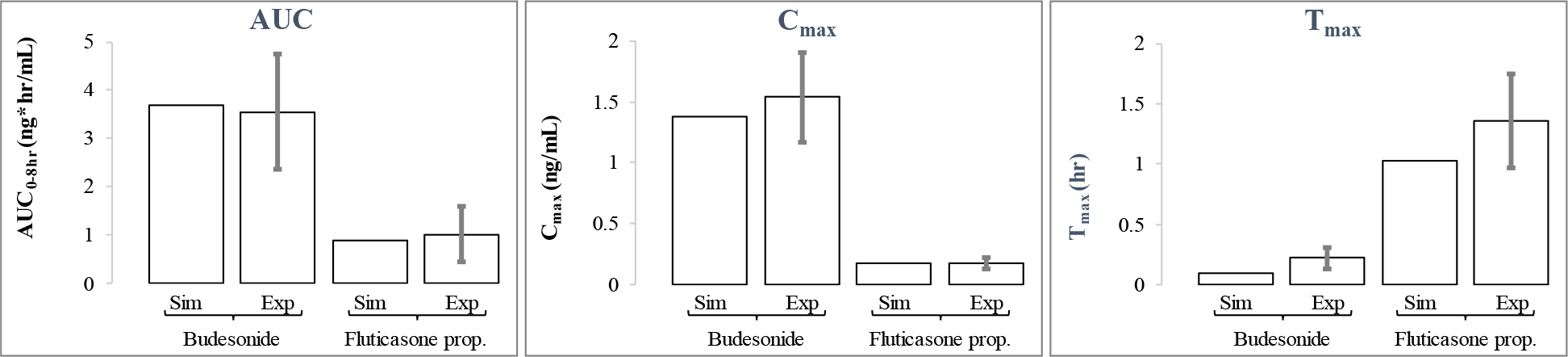
The simulated parameter comparison. The pharmacokinetics parameters comparison between simulation (Sim) data and average experimental (Exp) data with standard deviation bars. Experimental data points are calculated from digitalized raw data from multiple references.[85–89]

**Table 3.** Comparison of predicted budesonide pharmacokinetics parameters with average clinical data calculated from digitized raw data using absolute value of the difference between the two values.

| **Data** | **AUC<sub>0-8hr</sub> (ng\*hr/mL)** | **C<sub>max</sub> (ng/mL)** | **T<sub>max</sub> (hr)** |
| --- | --- | --- | --- |
| **Harrison et al. (2003)[85]** | 5.389 | 2.027 | 0.077187 |
| **Mortimer et al. (2007)[90]** | 2.392 | 1.67 | 0.21 |
| **Thorsson et al. (1994)[88]** | 2.923 | 1.507 | 0.3 |
| **<sup>a</sup>Thorsson et al. (1994)[88]** | 3.552 | 1.507 | 0.3 |
| **<sup>a</sup>Thorsson et al. (2001)[89]** | 4.52 | 1.636 | 0.28 |
| **Mollmann et al. (2001)[86]** | 2.53 | 0.9 | 0.17 |
| **Avg experimental data** | 3.551 | 1.541 | 0.222864 |
| **SD of experimental data** | 1.192 | 0.367 | 0.088709 |
| **Simulation** | 3.703 | 1.38 | 0.1 |

<sup>a</sup>Two data points from Thorsson et al. (1994) are from two different datasets in the same article. Also, the reported referenced experimental parameters are calculated from normalized pharmacokinetic plots shown in Fig 6.

**Table 4.** Comparison of predicted fluticasone propionate pharmacokinetics parameters with average clinical data calculated from digitized raw data.

| **Data** | **AUC<sub>0-8hr</sub> (ng\*hr/mL)** | **C<sub>max</sub> (ng/mL)** | **T<sub>max</sub> (hr)** |
| --- | --- | --- | --- |
| **Harrison et al. (2003)[85]** | 0.712 | 0.13 | 1.971477 |
| **Vurtikullird et al. (2016)[91]** | 0.71 | 0.221 | 1 |
| **Gillespie et al. (2015)[92]** | 1.684 | 0.224 | 1.1 |
| **Mortimer et al. (2007) [90]** | 0.4 | 0.12 | 1.21 |
| **Mollmann et al. (2001)[86]** | 1.58 | 0.188 | 1.5 |
| **Avg experimental data** | 1.017 | 0.177 | 1.356295 |
| **SD of experimental data** | 0.577 | 0.0493 | 0.391515 |
| **Simulation** | 0.89 | 0.181 | 1.033333 |

The reported referenced experimental parameters are calculated from normalized pharmacokinetic plots shown in Fig 6.

Fig 8 shows the pulmonary tissue retention profile of the two drugs through simulations.

**Fig 8.**
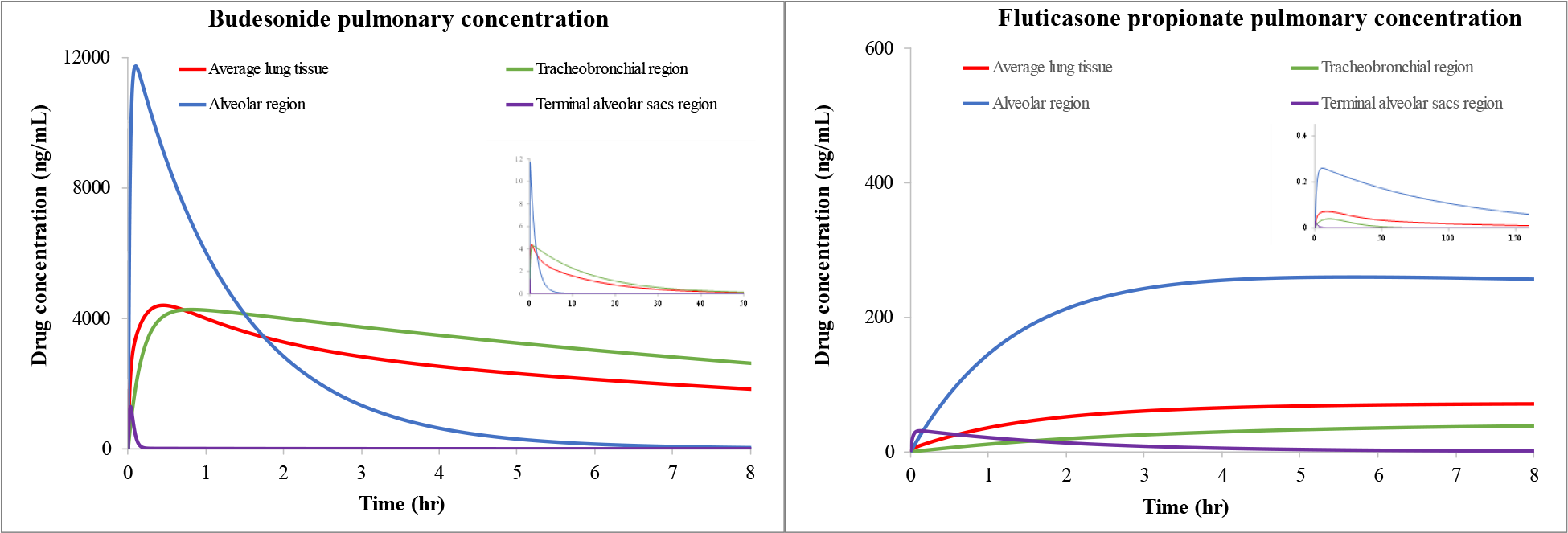
Pulmonary concentration-time profiles. Predicted average pulmonary concentration- time profiles in three different regions of the lung tissue as well as in total lung tissue, after 1 mg of drug inhalation of budesonide (left) and fluticasone propionate (right). The insert shows a larger simulation period of 50 hrs (for budesonide) and 150 hrs (for fluticasone propionate).

The provided values are for three different tissue regions (tracheobronchial, alveolar, and terminal alveolar sacs) along with the average of whole lung tissue, i.e., the average combination of three regions. Overall, it was observed that: i) both drugs stay in lung tissue (based on the concentration values) far longer than in the systemic blood (which is presented in Fig 6), ii) fluticasone propionate is retained in the lung significantly longer than budesonide, perhaps because of lower solubility and higher lipophilicity, iii) among different regions of the lung, Cmax is highest in alveolar region > tracheobronchial region > terminal alveolar sacs region, whereas

T1/2 (half life, time taken for Cmax to drop in half) is highest in tracheobronchial region (∼10bud and 30FP hrs) > alveolar region (∼1bud and 80FP hrs) > terminal alveolar sacs region (∼10bud and 100FP minutes), for both drugs.

### Parameter sensitivity

To investigate model sensitivity, some of the key model parameters were systematically varied. The optimal parameters that were used to obtain the concentration plots, shown in Fig 6 and Fig 8, were used as the baseline. The 12 individual parameters (except systemic clearance and logP) were varied from the base value by substituting high and low values (by increasing or decreasing by a factor of 2) into the model, while holding all other parameters constant. The parameters of systemic clearance and logP create physiologically unrealistic values if increased or decreased by a factor of 2, hence we created hypothetical upper and lower bounds for them to test sensitivity. Here systemic clearance was varied as 800 mL/min and 1800 mL/min for the lower and upper bounds, and logP was varied as increased or decreased by a factor of 1.5. The outcomes of parameter effects were quantified by comparing AUC0-8hr as shown in Fig 9-10. The individual AUC plots of parameter variations are shown in Fig 11-12.

**Fig 9.**
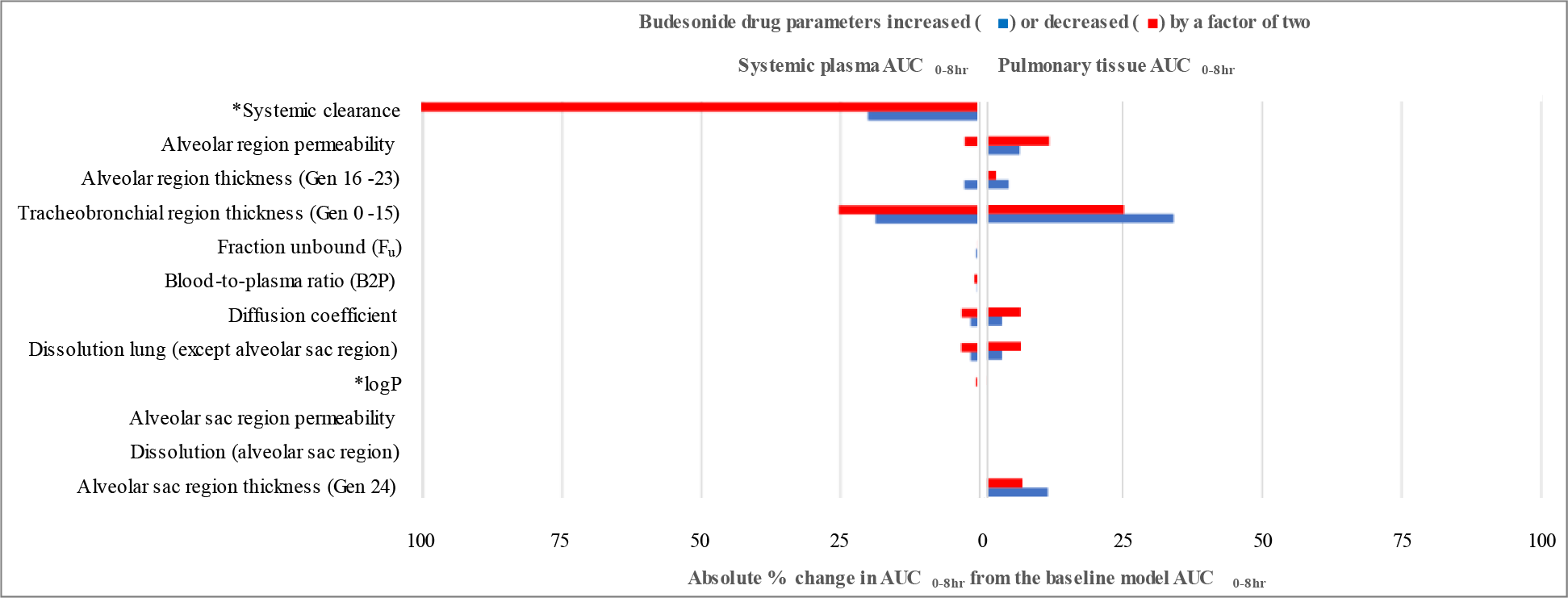
Sensitivity analysis of the input parameters (Budesonide). The sensitivity analysis is shown for the drug physicochemical and lung physiology parameters for budesonide in terms of the absolute percentage change in AUC0-8hr change from baseline systemic (left) and pulmonary tissue (right). The larger bars imply a stronger impact of the varied parameter on the respective pharmacokinetics outcome.

**Fig 10.**
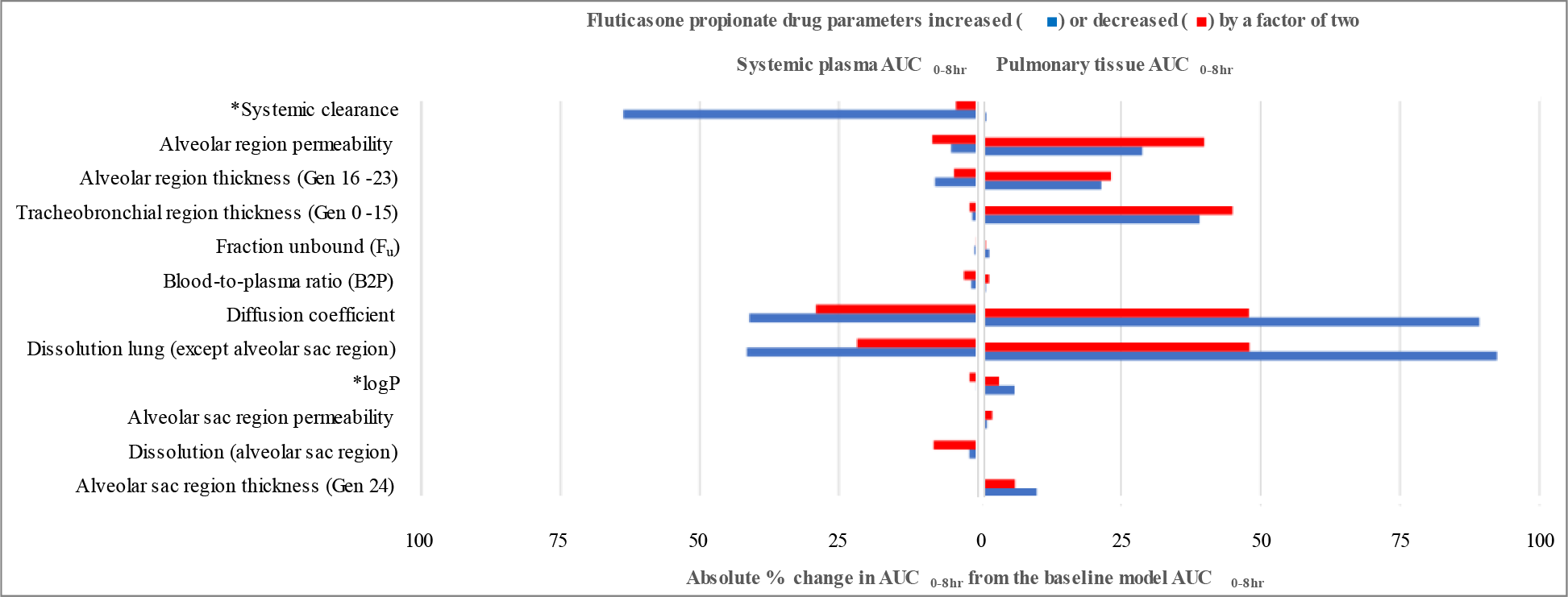
Sensitivity analysis of the input parameters (Fluticasone propionate). The sensitivity analysis is shown for the drug physicochemical and lung physiology parameters for fluticasone propionate in terms of the absolute percentage change in AUC0-8hr change from baseline systemic (left) and pulmonary tissue (right). The larger bars imply a stronger impact of the varied parameter on the respective pharmacokinetics outcome.

**Fig 11.**
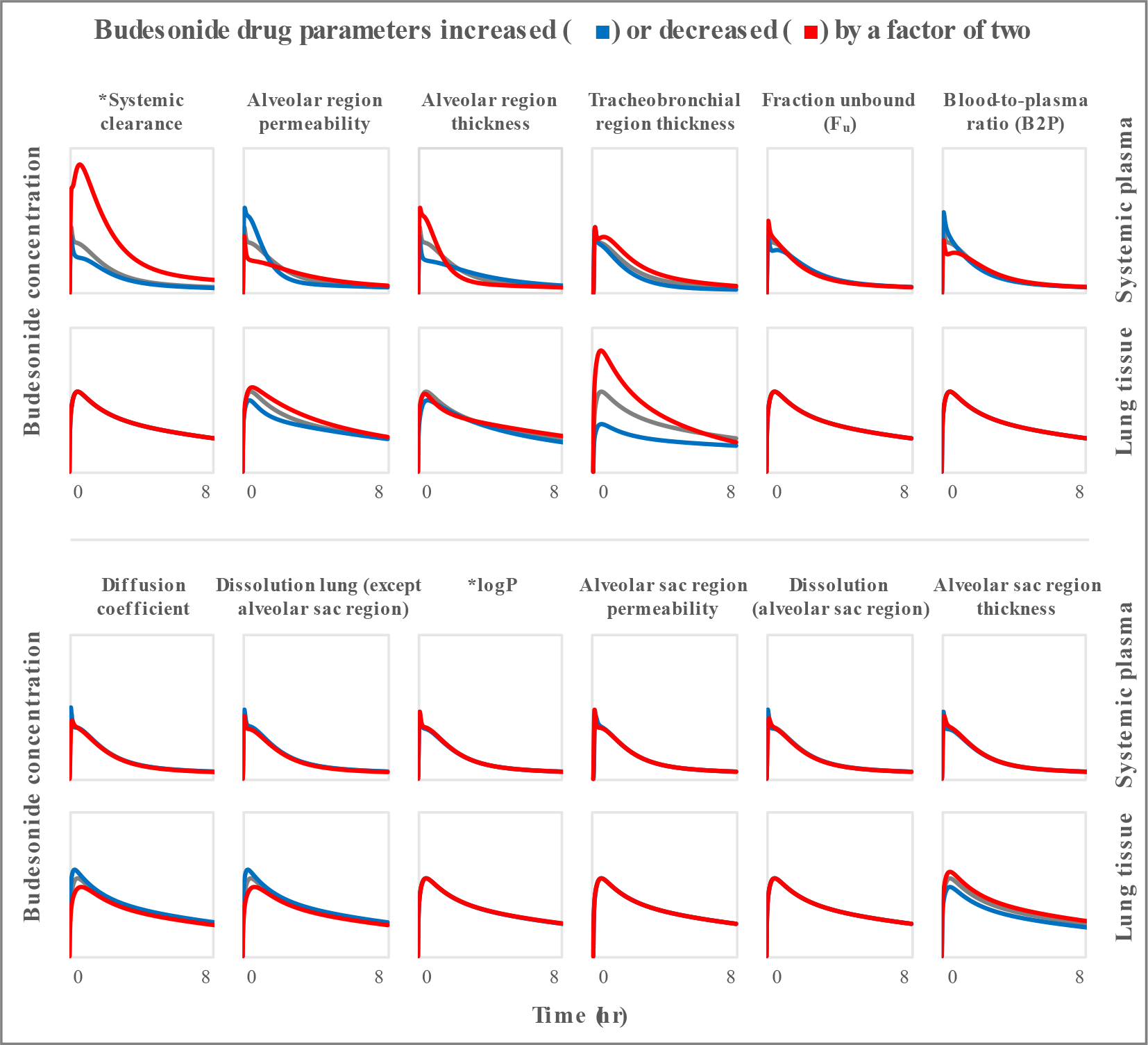
Parameter effects on drug concentration-time plots (Budesonide). Sensitivity analysis of drug physicochemical and lung physiology parameters for budesonide in terms of changes in drug concentrations as functions of time compared to baseline (gray line). The graphs in the upper row show systemic concentration and the lower row shows lung tissue concentration. Blue lines show parameters increased by a factor of two and the red line shows parameters decreased by a factor of two, with respect to baseline values.

**Fig 12.**
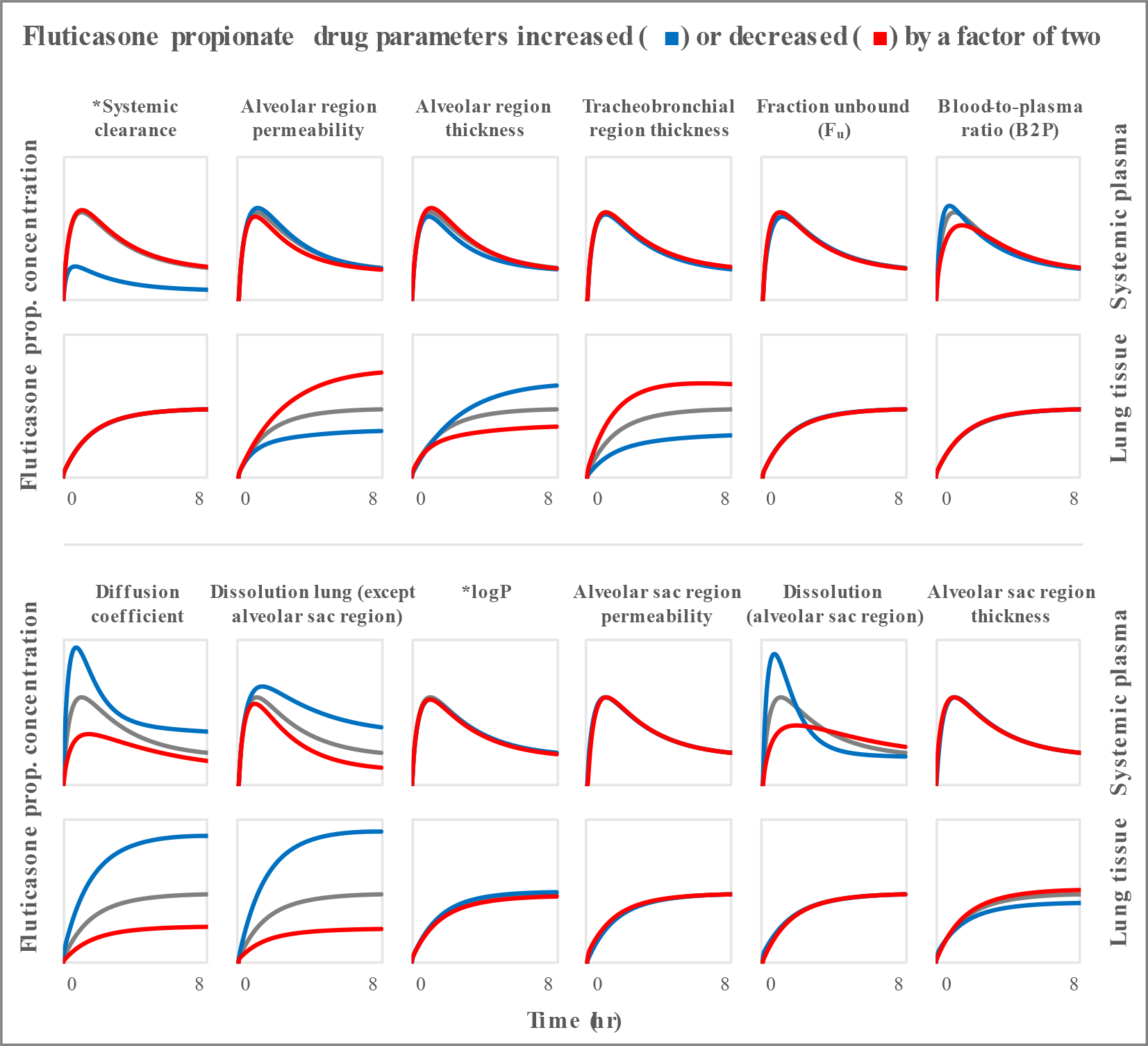
Parameter effects on drug concentration-time plots (Fluticasone propionate). Sensitivity analysis of drug physicochemical and lung physiology parameters for fluticasone propionate in terms of changes in drug concentrations as functions of time compared to baseline (gray line). The graphs in the upper row show systemic concentration and the lower row shows lung tissue concentration. Blue lines show parameters increased by a factor of 2 and the red line shows parameters decreased by a factor of 2, with respect to baseline values.

*For clarity, we have shown the axis cut-off of only 0-100% in these plots. The actual value of parameter “Systemic clearance” in systemic plasma AUC0-8hr plot above is 130%. As explained in Section *Parameter sensitivity*, the systemic clearance and logP parameters were varied differently than by a factor of two.

*As explained in Section *Parameter sensitivity*, the systemic clearance and logP parameters were varied differently than by a factor of two.

*As explained in Section *Parameter sensitivity*, the systemic clearance and logP parameters were varied differently than by a factor of two.

*As explained in Section Parameter sensitivity, the systemic clearance and logP parameters were varied differently than by a factor of two.

Overall, in determining the budesonide systemic drug concentration, the model was most sensitive to changes in systemic clearance and tracheobronchial barrier thicknesses. For budesonide lung tissue concentration, the model was most sensitive to changes in all the barrier thicknesses and drugs permeability.

For fluticasone propionate systemic drug concentration, the model was most sensitive to changes in systemic clearance, dissolution in lung fluid, and diffusion coefficient. For fluticasone propionate lung tissue concentration, the model was most sensitive to changes in all the barrier thicknesses and drugs permeability, dissolution in lung fluids, and diffusion coefficient.

## Discussion

Predictive tools for inhalation drug modeling have been published since the 1980s. The majority of these tools used compartmental modeling techniques that do not capture the complex 3D heterogeneity of human lungs and often involve the use of non-physiological parameters (for instance, simplified one-step drug translocation from the mucous to the plasma or not accounting for the regional barrier thicknesses). Since the site of action of these drugs is the lung tissue as a whole or specific lung regions that determine efficacy, predicting systemic concentration alone cannot be used to make predictions of any other events that happen in the lung tissue.

Unfortunately, systemic concentration is the only measurable outcome that can be validated with certainty in inhalation modeling, and multiple such in vivo (clinical) experimental datasets are available for different inhalatory drug types. The other outcome of predicted drug concentration in lung tissue is much more challenging and very few human studies have been published, with analysis limited to samples collected from bronchial biopsies (lavage or brushing).[30, 93] These, however, are limited to only providing information of the top epithelial layer mixed with mucosa and do not reflect the true drug concentration in the lung tissue itself. On the other hand, using pre-clinical animals models creates different types of challenges and uncertainties, such as: i) many common inhalers (DPIs and some metered dose inhalers) require breath actuation while most animals are nose-breathers, ii) species-specific heterogeneity in lung anatomy, and iii) different types of drug clearance mechanisms in animals as compared to humans.[94]

Hence, to gain a sound understanding of the features involved in the inhaled drug journey, the goal of this work was to develop and validate a mechanistic pulmonary PBPK model that can capture most of the relevant physiology and biophysics involved with the inhaled drug pathway (Fig 1). Model inputs include the breathing profile and drug PSD, employ all the relevant step-wise processes - deposition, dissolution, transport, and clearance, and the model provides final outcomes of drug concentration in systemic blood and different regions of the pulmonary tissue starting from throat-to-alveolar sacs. This outcome was compared to the clinical systemic pharmacokinetics data for budesonide and fluticasone propionate. Finally, a sensitivity analysis was performed to determine the most impactful physicochemical properties of drugs, formulation, and human physiological parameters for potential optimization to achieve a high lung selectivity and efficacy.

Additionally, this is the first instance of using a full-scale 3D lung model in a Q3D CFD framework to model the deposition, transport, and absorption of drugs in human lungs. The Q3D model is a simplified version of the 3D model, where the realistic 3D geometry is decomposed into a series of cylinders.[28] Such a geometry is well suited to model tubular structures like lungs as shown in Fig 3 and blood vessels,. The main advantage of using the Q3D approach to model drug absorption is that mucociliary transport of the undissolved and dissolved drug in the mucosa may be modeled with much greater precision than with a compartmental approach. It is possible that this enhanced precision will allow the PBPK model to simultaneously capture pulmonary and gastrointestinal tract absorptions with greater accuracy as discussed in a recent review of *in silico*methods for generic orally inhaled drug products.[13] In comparison, most other published studies have used simplified whole-lung dosimetry codes to predict particle deposition in the respiratory tract.[95, 96] The outcomes of these codes were used as inputs in further downstream modeling of drug pharmacokinetics.[21] Since the analytical/empirical equations in these codes were primarily designed for a bend, rather than a bifurcation, which changes the velocity flow path - they may not truly capture the deposition profile of inhaled lungs.[97]

### ICS simulations

Of the two different ICSs tested in the presented framework, budesonide has relatively high aqueous solubility (16-28 µg/mL in water and 470 µg/mL in 0.5% SDS that mimics some degree of mucosa/surfactant effects), whereas fluticasone propionate is practically insoluble (>0.1 µg/mL) in water and sparingly soluble in SDS. In addition, the difference in lipophilicity between the two drugs affects the dissolution rate of the drug that is deposited in the mucosa. This low solubility has a three-fold effect on fluticasone propionate pharmacokinetics. First, the prolonged presence of deposited, undissolved particles of fluticasone propionate in the mucosa exposes the drug for longer clearance mechanisms by MCC.[23, 31] With this mechanism, the drug further travels from throat-to-mouth-to-gut. In the gut, the final drug absorption in the systemic circulation is determined by the bioavailability fraction of the drug as well as the other liver/kidney clearance mechanisms. Second, the prolonged presence of the drug in mucosa is reflected in slow and extended absorption/transport of the drug in lung tissue barriers, and hence, it can be expected that pulmonary tissue pharmacokinetics of drug will be observed for many hours/days. However, no clear experimental data are available to support this. Limited proxy experiments have been published in human bronchial brush samples that observed the fluticasone propionate concentration in samples even after 18 hours post inhalation.[93] Third, since the bioavailability of fluticasone propionate is less than 1% in the gut,[98] the systemic drug contribution back to the pulmonary region (through pulmonary circulation) will be minimal. In comparison, budesonide has a gut bioavailability of ∼10%, and hence, it is expected that the fraction of drug that gets absorbed in systemic circulation through the gut will travel back to the pulmonary region. However, this gut absorbed fraction will only have minor effect on plasma pharmacokinetics. For examples, in our simulations (Table 2), 40% of the total drug is deposited in mouth-throat region. This means that only 4% of the total systemic drug contribution comes from gut absorbed fraction (10% bioavailable fraction of the 40% swallowed fraction from mouth-throat).

### Bi-phasic response

Along with efficiently simulating the pharmacokinetic responses of inhaled ICS drugs, the above-described model can also be used to provide mechanistic insights into phasic responses. For examples, the in vivo pharmacokinetic studies of Mollmann et al.[86] and Harrison et al.[85] (Fig 6) has shown a delayed second peak after 10-20 minutes (bi-phasic response) in budesonide pharmacokinetics. This has also been captured in the presented simulation results. We hypothesize that this could be due to the difference in absorption efficiency of different lung regions, i.e., the deposited drug can get absorbed much faster in the terminal alveolar sacs or alveolar region due to their thin barriers compared to the thick barriers of the conducting region. To identify this regional contribution, we systematically switched off (blocked) one region at a time and observed the resulting pharmacokinetic profile of inhaled budesonide while keeping everything else same.

This analysis (Fig 13) shows that: 1) gut absorption has minimal effect on systemic concentration (gut block vs original simulation), 2) the early peak (Cmax) is due to the fast absorption from terminal alveolar sacs region as well as alveolar region (tracheobronchial region block vs original simulation), 3) the sharp peak is still present after alveolar region block, and absent when terminal alveolar sacs region is blocked, implying that terminal sacs are primarily responsible for rapid sharp peak of drug concentration after inhalation. Additionally, the bi- phasic response may be expected based on the PSD profile of inhaled drugs. For example, it is possible that post-inhalation a fraction of the smallest size drug particles that directly reach the terminal alveolar sacs region rapidly permeate through the thin air-blood barrier, especially if the drug solubility is high as in the case of budesonide, thereby causing a rapid and early spike in systemic blood concentration. Naturally, this also implies that once most of this deposited drug is quickly absorbed from this region, a sudden drop in drug concentration will be observed before other alveolar region absorption starts contributing to the systemic concentration, causing the second peak. However, this mechanistic hypothesis has not yet been explored in any experimental studies. Nonetheless, due to such differences in the properties of these two drugs, one can expect a short Tmax and a much faster rise in drug concentration in the blood (Cmax) for budesonide in comparison with fluticasone propionate. This has been observed in multiple in vivo pharmacokinetics studies and well matched in presented simulations as shown in Fig 6.

**Fig 13.**
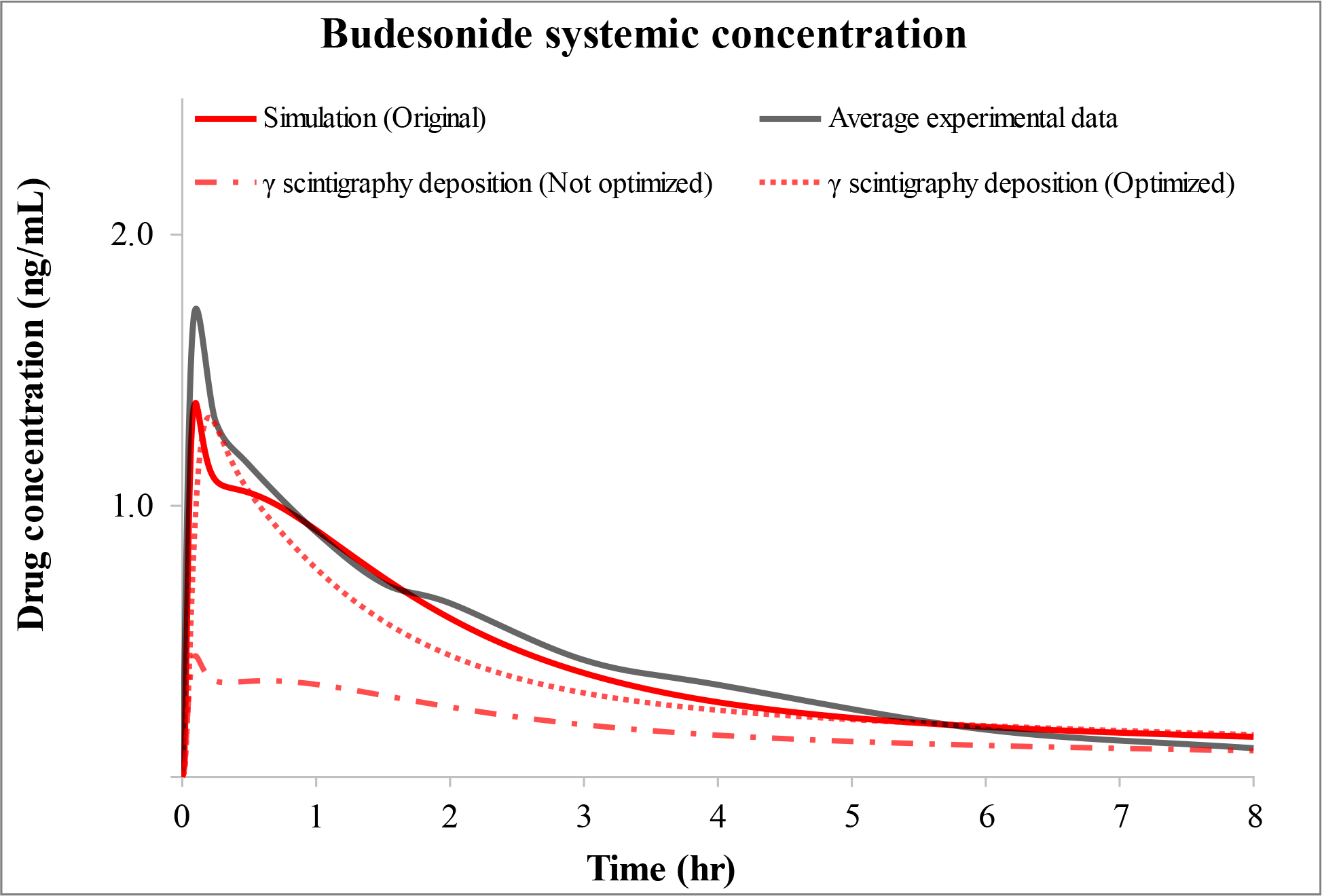
Regional contribution in overall systemic concentration (Budesonide). The results shown for the plasma (systemic) concentration-time profiles after administration of 1 mg of inhaled budesonide. Original simulations results are compared with switching off (blocking) one region at a time, while keeping everything else same.

### Impact of change in regional drug deposition

As reported in the Drug deposition Section (Results), the Q3D-predicted total lung deposition fraction for budesonide (47% of the metered dose), falls outside of the experimental range of 9.4 – 41% (median 32.1%) described by in vivo *γ* scintigraphy studies conducted by Newman et al. [29] This raised the question: Had Q3D predicted the same deposition profile as calculated by Newman et al. [29], how would that change the predicted systemic pharmacokinetic outcome? To address this, the budesonide pharmacokinetic simulations were repeated using the input deposition fraction values provided by Newman et al. [29] Two approaches were used: (i) the systemic drug concentration was computed while retaining the previously calibrated parameters based on the Q3D deposition fraction shown in Table 1, and (ii) the systemic drug concentration was computed by recalibration of these parameters to match the average experimental (clinical) pharmacokinetic profile. The first approach is to gain insight into the impact regional deposition can have on the systemic pharmacokinetic profile in the absence of other parameter adjustments, while the latter is to ascertain the impact that different regional deposition predictions would have during the usual course of model development. As shown in Fig 14 and Table 5, differences in drug deposition fractions had easily visible impacts on predicted plasma concentration of the drug. The 14.9% decrease in deposited drug (47% to 32.1%) resulted in a predicted decrease for AUC0-8hr on a relative basis of ∼60% and ∼20% using the first and second approaches, respectively. The predicted relative decrease in Cmax was only about 4% using the second approach but using the first approach the predicted relative decrease was about 68%. The predicted change in Tmax was minor.

**Fig 14.**
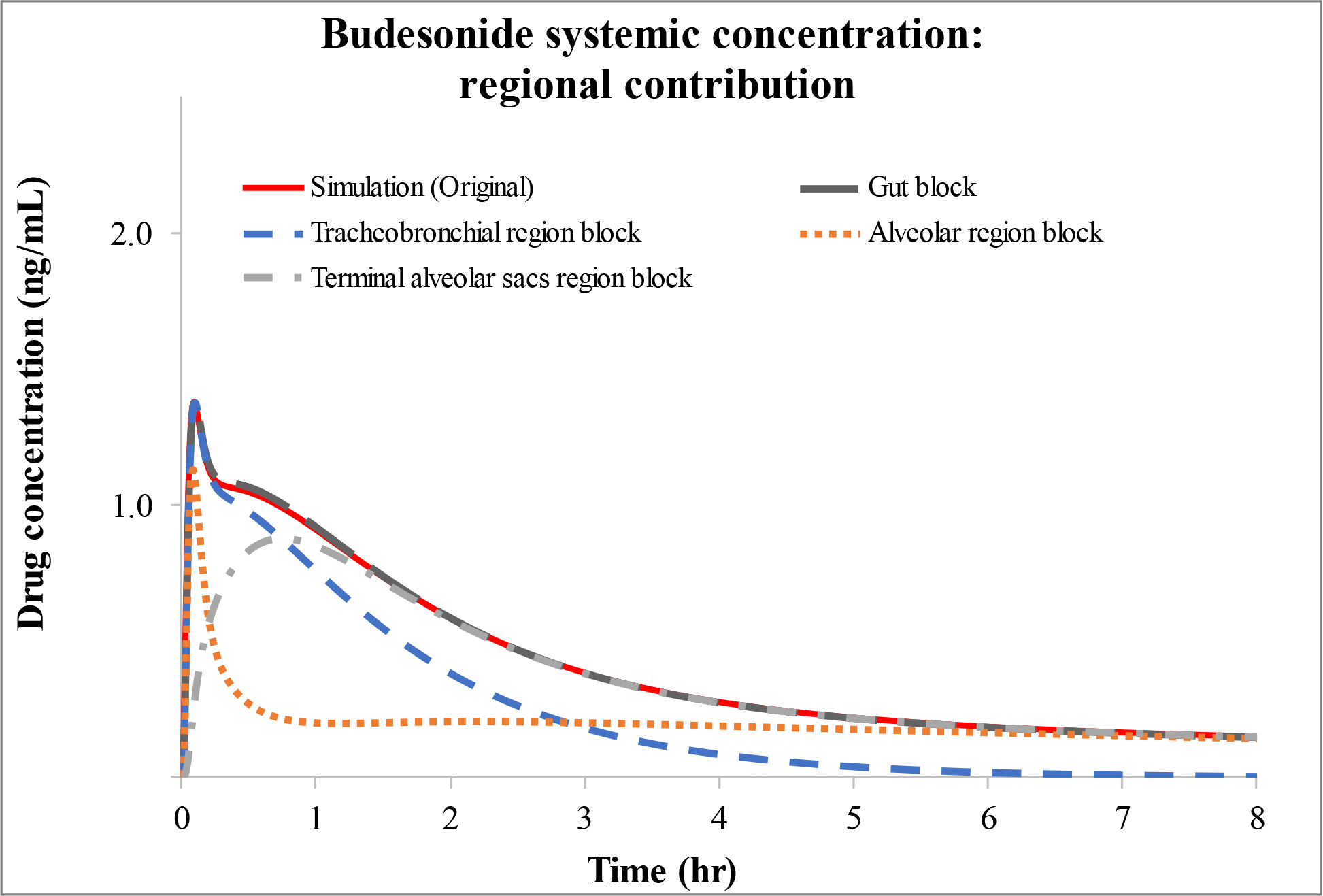
Plasma (systemic) concentration-time profiles after administration of 1 mg of inhaled budesonide. The plots of Average experimental data and Simulation (Original) predictions are the same as in Fig 6 and are provided for the sake of comparison. In one case, the Newman et al. [29] *γ* scintigraphy data are used as inputs while keeping everything (drug and barrier parameters) same as Simulation (Original), while in the other case these parameters were optimized to get the best fit with respect to average experimental data.

**Table 5.**
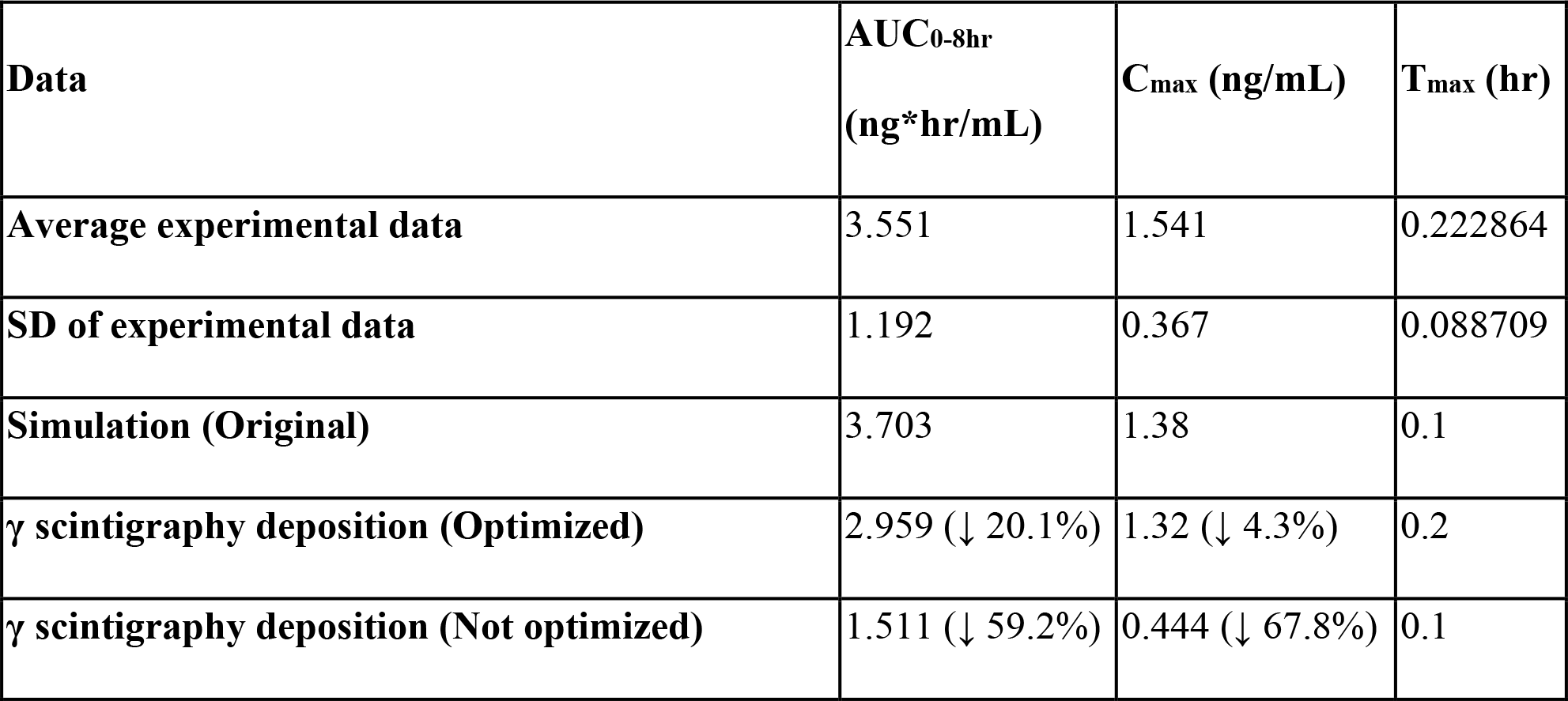
Comparison of simulated budesonide pharmacokinetics parameters with average clinical data calculated from digitized raw data.

### Lung tissue concentration

For pulmonary tissue concentration, the goal of this study is to look at the *trend* of drug pharmacokinetics in different regions of the lung and not the actual values of tissue concentration *per se* since it cannot be validated by experimental analysis. As shown in Fig 8, the first main observation of the simulated outcomes is that the tissue affinity (lung retention profile) for fluticasone propionate is much larger compared to budesonide. Mechanistically, this is generally positively correlated to the drug’s lipophilicity represented by logP (budesonide = 2.3, fluticasone propionate = 3.7) and the dissolution rate (which is much lower for fluticasone propionate) of the drug particles deposited in the lung lumen. Previous *in vitro* studies on dissolution rates of these two drugs have shown that while budesonide particles were dissolved within 6 minutes, fluticasone propionate required at least 6-8 hours.[23, 31] Others have shown that only 6-7% of fluticasone propionate deposited on different human lung cells was absorbed through the cell monolayer during 4 hours, whereas 10 times (60-70%) more budesonide was transported through the same cell line in the same time period.[99–102] The prolonged presence of fluticasone propionate in the airway lumen and slow absorption in lung tissue is also reflected in the much longer time for systemic absorption of fluticasone propionate than that of budesonide (Fig 6). The second main observation is that both drugs showed higher concentrations in the alveolar region compared to the tracheobronchial region. Experimentally, this trend was also observed by Himstedt et al. for fluticasone propionate and other respiratory drugs in animal (rat) models that found a six-fold higher drug affinity for the alveolar parenchyma than the trachea.[103] However, that study used intravenous administration of drugs in animals and may not reveal the true dynamics of inhalation administration. The third main observation is that the average tissue concentration of budesonide can be observed in lung tissue for up to 40 hours compared to an even longer time period for fluticasone propionate (150+ hours, Fig 8). Surprisingly, budesonide stays in tracheobronchial region for a longer time compared to alveolar and alveolar sacs regions due to the slower translocation across the thicker tracheobronchial barriers after faster dissolution but lower lipophilicity. In comparison, the fluticasone propionate stays in alveolar region for much longer than tracheobronchial region due to very slow dissolution rate and lack of MCC in alveolar region. However, no experimental support can be found in published literature to support such long-term region-specific response of these simulated drugs.

### Model parameter sensitivity

To investigate model sensitivity, we systematically varied some of the physicochemical and physiological parameters (Fig 9-12). The overall analysis showed a clear difference in parameter sensitivity between systemic and pulmonary outcomes, as well as, between the two ICSs. Among plasma-related parameters, systemic clearance had a significant effect on both drugs, while B2P and fu did not induce much change in AUC0-8hr values in either of the drugs.

For systemic clearance, it is important to note that parameter change by a factor of two created unphysiological values and hence we picked 800-1800 mL/min as low and high values to test. Since our baseline itself is ∼1600 mL/min for budesonide and ∼850 mL/min for fluticasone propionate, it did create some discrepancy in looking at the low and high range effects.

Nonetheless, the rate of drug clearance in the blood is one of the most significant parameters to determine systemic AUC.

Among physicochemical parameters, diffusion coefficient and dissolution (both determined by solubility values) changes had minor effects on budesonide pharmacokinetics, both in systemic and pulmonary tissue. The two-fold increase and decrease in these parameters only caused a ∼5-10% change in AUC0-8hr compared to baseline. In comparison, for fluticasone propionate, these parameters induced up to 20-40% change in systemic and 50-90% change in lung tissue concentration. It is possible that the changes are higher in fluticasone propionate because the starting (baseline) value itself is significantly low (i.e., practically insoluble) and any minor change significantly induces higher dissolution of the deposited drug in the mucosa. For budesonide the solubility is already optimal (very soluble) at baseline, and hence, only minor changes are observed by the change in these parameters. On other hand, as expected, the effect of systemic clearance induced minimal effects in pulmonary tissue concentration, whereas the tissue barrier thicknesses were more significant for both the drugs, where changes to the AUC of 5-30% in budesonide and 22-48% in fluticasone propionate were predicted. For budesonide, which already has a large solubility coefficient and therefore differences in this parameter do not play much of role in how fast the drug gets transported into the tissue, it is only the thickness of tracheobronchial barrier which significantly influence the speed and amount with which drug permeates into the blood. For much smaller thicknesses of alveolar and terminal alveolar sacs regions, the translocation from these sections is already rapid due to high solubility of budesonide. In comparison, solubility coefficient and diffusion are the main drivers for fluticasone propionate transport into the lung tissue barriers. Here the solubility coefficient is very small and hence a small change to that will result in a great change to the translocating rate into the tissue. Similarly, the solubility equation has the diffusion coefficient as a pre-multiplier, hence, the effect of the diffusion coefficient is also significant in determining fluticasone propionates AUC changes in sensitivity analysis.

Overall, most physiological outcomes have nonlinear dependency on any particular parameter, and the net change in AUC or transport rate, etc., is likely to be a complex interplay of the individual parameter fluxes. Hence, the goal for these type of fast-running ‘what-if’ scenarios was to: i) explore what type of drug parameters can be explored *a priori* before the experimental studies (such as formulation design) to increase drug efficacy and reduce systemic toxicity, ii) identify parameter specific, or combinatory effects of parameters, to explore lung selectivity index (ratio between pulmonary and systemic exposure ratio) of inhaled drugs, and iii) to help other modelers in optimizing the lung barrier models for related studies.

## Limitations

As discussed above, the primary limitation for validating lung pharmacokinetics models is that there is a lack of pulmonary tissue concentration data, so it is generally not possible to validate the model against the true metrics of interest. Until this issue is addressed with experimental support, lung pharmacokinetics model validation will likely be limited to comparison with systemic drug concentration values, which does not ensure that site of action tissue concentration predictions are accurate. The closest available comparison is between predicted and experimentally observed values of regional deposition, where the consequences of potential differences in regional deposition was explored as shown in Table 5 and Fig 13. However, regional absorption may be different than regional deposition if absorption is dissolution- or permeability-limited. Other potential limitations in our current framework are a lack of device specific effects (such as single actuation content and carrier effects for DPIs and plume geometry and spray pattern for MDIs),[13, 104] a lack of other clearance mechanisms (such as drug phagocytosis by alveolar macrophages and cleared by transport to the lung-draining lymph nodes),[105, 106] and lung region-specific involvement of metabolic and transported enzymes and proteins that may modulate the lung retention and bioavailability of some drugs.[107] Hence, overall it is possible that the lung tissue concentration of inhaled drugs may be overpredicted in absence of these modules in the model framework. A goal in future versions of this model is to resolve these limitations and thereby improve the prediction process. Further, as clinical trial data of systemic pharmacokinetics often involves a mixed population (male and female participants), an equivalent female lung model should also be part of OIDP prediction framework.

In conclusion, the presented model is a comprehensive fully mechanistic and physiologically realistic computational framework that captures multiple processes that are essential to describe the fate of inhaled drug kinetics. The work also highlights the importance of drug parameters and physiologic differences between different regions of lung tissues and their impact on systemic as well as lung retention profile. The expected applications are improvements in the qualitative and quantitative understanding of inhaled drug behavior, optimization of drugs and formulations for improved and targeted efficacy, and to aid in the design of clinical trials.

## Supporting information

**S1 Appendix.** Further details of model’s aerosol transport and deposition equations along with the mesh independence analysis.

## Acknowledgments

Views expressed in this work do not necessarily reflect the official policies of the U.S. Food and Drug Administration (FDA); nor does any mention of trade names, commercial practices, or organization imply endorsement by the U.S. Government. The authors gratefully acknowledge the guidance and technical feedback of FDA-CDER team.

## Author Contributions

Narender Singh

ROLES: Conceptualization, Data curation, Formal analysis, Investigation, Methodology, Resources, Model Software, Visualization, Writing – original draft, Writing – review & editing AFFILIATION: CFD Research, Huntsville, Alabama, United States of America

Ravi Kannan

ROLES: Methodology, Resources, Model Software, Visualization, Writing – original draft, Writing – review & editing

AFFILIATION: CFD Research, Huntsville, Alabama, United States of America

Ryan Arey

ROLES: Methodology, Resources, Model Software

AFFILIATION: CFD Research, Huntsville, Alabama, United States of America

Ross Walenga

ROLES: Conceptualization, Resources, Writing – review & editing

AFFILIATION: Division of Quantitative Methods and Modeling, Office of Research and Standards, Office of Generic Drugs, Center for Drug Evaluation and Research, U.S. Food and Drug Administration, Silver Spring, Maryland, United States of America

Andrew Babiskin

ROLES: Conceptualization, Resources, Writing – review & editing

AFFILIATION: Division of Quantitative Methods and Modeling, Office of Research and Standards, Office of Generic Drugs, Center for Drug Evaluation and Research, U.S. Food and Drug Administration, Silver Spring, Maryland, United States of America

Andrzej Przekwas

ROLES: Conceptualization, Methodology, Resources, Writing – review & editing AFFILIATION: CFD Research, Huntsville, Alabama, United States of America

## References

1. Report (2018) GBD 2017: a fragile world. Lancet 392: 1683.

2. Chow AH, Tong HH, Chattopadhyay P, Shekunov BY (2007) Particle engineering for pulmonary drug delivery. Pharm Res 24: 411–37.

3. Labiris NR, Dolovich MB (2003) Pulmonary drug delivery. Part I: physiological factors affecting therapeutic effectiveness of aerosolized medications. Br J Clin Pharmacol 56: 588–99.

4. Cooper AE, Ferguson D, Grime K (2012) Optimisation of DMPK by the inhaled route: challenges and approaches. Curr Drug Metab 13: 457–73.

5. Goel A, Baboota S, Sahni JK, Ali J (2013) Exploring targeted pulmonary delivery for treatment of lung cancer. Int J Pharm Investig 3: 8–14.

6. Hess DR (2008) Aerosol delivery devices in the treatment of asthma. Respir Care 53: 699–723; discussion 723-5.

7. van Noord JA, Smeets JJ, Maesen FP (1998) A comparison of the onset of action of salbutamol and formoterol in reversing methacholine-induced bronchoconstriction. Respir Med 92: 1346–51.

8. Borghardt JM, Kloft C, Sharma A (2018) Inhaled Therapy in Respiratory Disease: The Complex Interplay of Pulmonary Kinetic Processes. Can Respir J 2018: 2732017.

9. Lipworth BJ (1999) Systemic adverse effects of inhaled corticosteroid therapy: A systematic review and meta-analysis. Arch Intern Med 159: 941–55.

10. Patton JS, Byron PR (2007) Inhaling medicines: delivering drugs to the body through the lungs. Nat Rev Drug Discov 6: 67–74.

11. Ibrahim M, Verma R, Garcia-Contreras L (2015) Inhalation drug delivery devices: technology update. Med Devices (Auckl) 8: 131–9.

12. Zhong H, Chan G, Hu Y, Hu H, Ouyang D (2018) A Comprehensive Map of FDA-Approved Pharmaceutical Products. Pharmaceutics 10.

13. Walenga RL, Babiskin AH, Zhao L (2019) In Silico Methods for Development of Generic Drug-Device Combination Orally Inhaled Drug Products. CPT Pharmacometrics Syst Pharmacol 8: 359–70.

14. Weber B, Hochhaus G (2013) A pharmacokinetic simulation tool for inhaled corticosteroids. AAPS J 15: 159–71.

15. Boger E, Friden M (2019) Physiologically Based Pharmacokinetic/Pharmacodynamic Modeling Accurately Predicts the Better Bronchodilatory Effect of Inhaled Versus Oral Salbutamol Dosage Forms. J Aerosol Med Pulm Drug Deliv 32: 1–12.

16. Hendrickx R, Lamm Bergstrom E, Janzen DLI, Friden M, Eriksson U, et al. (2018) Translational model to predict pulmonary pharmacokinetics and efficacy in man for inhaled bronchodilators. CPT Pharmacometrics Syst Pharmacol 7: 147–57.

17. Gaz C, Cremona G, Panunzi S, Patterson B, De Gaetano A (2012) A geometrical approach to the PKPD modelling of inhaled bronchodilators. J Pharmacokinet Pharmacodyn 39: 415–28.

18. Caniga M, Cabal A, Mehta K, Ross DS, Gil MA, et al. (2016) Preclinical Experimental and Mathematical Approaches for Assessing Effective Doses of Inhaled Drugs, Using Mometasone to Support Human Dose Predictions. J Aerosol Med Pulm Drug Deliv 29: 362–77.

19. Hartung N, Borghardt JM (2020) A mechanistic framework for a priori pharmacokinetic predictions of orally inhaled drugs. PLoS Comput Biol 16: e1008466.

20. Chaudhuri SR, Lukacova V (2010) Simulating delivery of pulmonary (and intranasal) aerosolised drugs. ONdrugDelivery: 26–30.

21. Borghardt JM, Weber B, Staab A, Kloft C (2015) Pharmacometric Models for Characterizing the Pharmacokinetics of Orally Inhaled Drugs. AAPS J 17: 853–70.

22. Backman P, Arora S, Couet W, Forbes B, de Kruijf W, et al. (2018) Advances in experimental and mechanistic computational models to understand pulmonary exposure to inhaled drugs. Eur J Pharm Sci 113: 41–52.

23. Edsbacker S, Wollmer P, Selroos O, Borgstrom L, Olsson B, et al. (2008) Do airway clearance mechanisms influence the local and systemic effects of inhaled corticosteroids? Pulm Pharmacol Ther 21: 247–58.

24. Sakagami M, Kinoshita W, Sakon K, Makino Y (2003) Fractional contribution of lung, nasal and gastrointestinal absorption to the systemic level following nose-only aerosol exposure in rats: a case study of 3.7- micro m fluorescein aerosols. Arch Toxicol 77: 321–9.

25. Sturm R (2011) Age-dependence and intersubject variability of tracheobronchial particle clearance. Pneumon 24: 77–85.

26. Kannan R, Singh N, Przekwas A, Zhou X, Walenga R, et al. (2021) A Quasi-3D model of the whole lung: Airway extension to the tracheobronchial limit using the constrained constructive optimization and alveolar modelling, using a sac-trumpet model. J Comput Des Eng (ACCEPTED).

27. Kannan R, Chen ZJ, Singh N, Przekwas A, Delvadia R, et al. (2017) A quasi-3D wire approach to model pulmonary airflow in human airways. Int J Numer Method Biomed Eng 33.

28. Kannan R, Singh N, Przekwas AJ (2017) A compartment-quasi3D multiscale approach for drug absorption, transport, and retention in the human lungs. Int J Numer Method Biomed Eng.

29. Newman SP, Pitcairn GR, Hirst PH, Bacon RE, O’Keefe E, et al. (2000) Scintigraphic comparison of budesonide deposition from two dry powder inhalers. Eur Respir J 16: 178–83.

30. Dalby C, Polanowski T, Larsson T, Borgstrom L, Edsbacker S, et al. (2009) The bioavailability and airway clearance of the steroid component of budesonide/formoterol and salmeterol/fluticasone after inhaled administration in patients with COPD and healthy subjects: a randomized controlled trial. Respir Res 10: 104.

31. Johnson M (1996) Pharmacodynamics and pharmacokinetics of inhaled glucocorticoids. J Allergy Clin Immunol 97: 169–76.

32. Kannan R, Przekwas AJ, Singh N, Delvadia R, Tian G, et al. (2017) Pharmaceutical aerosols deposition patterns from a Dry Powder Inhaler: Euler Lagrangian prediction and validation. Med Eng Phys 42: 35–47.

33. Kannan R, Singh N, Przekwas AJ (2018) A quasi-3D compartmental multi-scale approach to detect and quantify diseased regional lung constriction using spirometry data. Int J Numer Method Biomed Eng 34.

34. Kannan R, Guo P, Przekwas AJ (2015) Particle transport in the human respiratory tract: formulation of a nodal inverse distance weighted Eulerian–Lagrangian transport and implementation of the Wind–Kessel algorithm for an oral delivery. Int J Numer Method Biomed Eng 32.

35. Miyawaki S, Choi S, Hoffman EA, Lin CL (2016) A 4DCT imaging-based breathing lung model with relative hysteresis. J Comput Phys 326: 76–90.

36. Rajaraman PK, Choi J, Hoffman EA, O’Shaughnessy PT, Choi S, et al. (2020) Transport and deposition of hygroscopic particles in asthmatic subjects with and without airway narrowing. J Aerosol Sci 146: 105581.

37. Karch R, Neumann F, Neumann M, Schreiner W (1999) A three-dimensional model for arterial tree representation, generated by constrained constructive optimization. Comput Biol Med 29: 19–38.

38. Pichelin M, Caillibotte G, Katz I, Martonen T (2012) Categorization of Lung Morphology Based on FRC and Height: Computer Simulations of Aerosol Deposition. Aerosol Sci Technol 46: 70–81.

39. Weibel ER (1991) Design of airways and blood vessels as branching trees. New York: Raven Press.

40. Mercer RR, Russell ML, Roggli VL, Crapo JD (1994) Cell number and distribution in human and rat airways. Am J Respir Cell Mol Biol 10: 613–24.

41. Frohlich E, Mercuri A, Wu S, Salar-Behzadi S (2016) Measurements of Deposition, Lung Surface Area and Lung Fluid for Simulation of Inhaled Compounds. Front Pharmacol 7: 181.

42. Pinkerton KE, Gehr P, Castañeda A, Crapo JD (2015) Chapter 9 - Architecture and Cellular Composition of the Air–Blood Tissue Barrier. In: Parent RA, editor. Comparative Biology of the Normal Lung (Second Edition). San Diego: Academic Press. pp. 105–17.

43. Yu JY, Rosania GR (2010) Cell-based multiscale computational modeling of small molecule absorption and retention in the lungs. Pharm Res 27: 457–67.

44. Olsson B, Bondesson E, Borgström L, Edsbäcker S, Eirefelt S, et al. (2011) Pulmonary Drug Metabolism, Clearance, and Absorption. In: Smyth HDC, Hickey AJ, editors. Controlled Pulmonary Drug Delivery. New York, NY: Springer New York. pp. 21–50.

45. Wauthoz N, Amighi K (2015) Formulation Strategies for Pulmonary Delivery of Poorly Soluble Drugs. In: Nokhodchi A, Martin GP, editors. Pulmonary Drug Delivery: Advances and Challenges: John Wiley & Sons, Ltd.

46. (1977) Ozone and Other Photochemical Oxidants. Washington, DC: The National Academies Press.

47. Tian G, Hindle M, Lee S, Longest PW (2015) Validating CFD Predictions of Pharmaceutical Aerosol Deposition with In Vivo Data. Pharm Res 32: 3170-87.

48. Hill LS, Slater AL (1998) A comparison of the performance of two modern multidose dry powder asthma inhalers. Respir Med 92: 105–10.

49. Daley-Yates PT (2015) Inhaled corticosteroids: potency, dose equivalence and therapeutic index. Br J Clin Pharmacol 80: 372–80.

50. Csizmadia F, Tsantili-Kakoulidou A, Panderi I, Darvas F (1997) Prediction of distribution coefficient from structure. 1. Estimation method. J Pharm Sci 86: 865–71.

51. Tetko IV, Tanchuk VY (2002) Application of associative neural networks for prediction of lipophilicity in ALOGPS 2.1 program. J Chem Inf Comput Sci 42: 1136–45.

52. Jones RM, Harrison A (2012) A new methodology for predicting human pharmacokinetics for inhaled drugs from oratracheal pharmacokinetic data in rats. Xenobiotica 42: 75–85.

53. Derendorf H, Hochhaus G, Meibohm B, Mollmann H, Barth J (1998) Pharmacokinetics and pharmacodynamics of inhaled corticosteroids. J Allergy Clin Immunol 101: S440–6.

54. Lombardo F, Berellini G, Obach RS (2018) Trend Analysis of a Database of Intravenous Pharmacokinetic Parameters in Humans for 1352 Drug Compounds. Drug Metab Dispos 46: 1466–77.

55. Szefler SJ (1999) Pharmacodynamics and pharmacokinetics of budesonide: a new nebulized corticosteroid. J Allergy Clin Immunol 104: 175–83.

56. Andersson P, Brattsand R, Dahlstrom K, Edsbacker S (1993) Oral availability of fluticasone propionate. Br J Clin Pharmacol 36: 135–6.

57. Boger E, Evans N, Chappell M, Lundqvist A, Ewing P, et al. (2016) Systems Pharmacology Approach for Prediction of Pulmonary and Systemic Pharmacokinetics and Receptor Occupancy of Inhaled Drugs. CPT Pharmacometrics Syst Pharmacol 5: 201–10.

58. Falcoz C, Oliver R, McDowall JE, Ventresca P, Bye A, et al. (2000) Bioavailability of orally administered micronised fluticasone propionate. Clin Pharmacokinet 39 Suppl 1: 9–15.

59. https://go.drugbank.com/drugs/DB01222.

60. https://www.accessdata.fda.gov/drugsatfda_docs/label/2009/021324s008lbl.pdf.

61. Winkler J, Hochhaus G, Derendorf H (2004) How the lung handles drugs: pharmacokinetics and pharmacodynamics of inhaled corticosteroids. Proc Am Thorac Soc 1: 356–63.

62. Odziomek M, Sosnowski TR, Gradoń L (2015) The Influence of Functional Carrier Particles (FCPs) on the Molecular Transport Rate Through the Reconstructed Bronchial Mucus: In Vitro Studies. Transport in Porous Media 106: 439-54.

63. Tashkin DP, Lipworth B, Brattsand R (2019) Benefit:Risk Profile of Budesonide in Obstructive Airways Disease. Drugs 79: 1757–75.

64. Nakowitsch S, Koller C, Seifert JM, Konig-Schuster M, Unger-Manhart N, et al. (2020) Saponin Micelles Lead to High Mucosal Permeation and In Vivo Efficacy of Solubilized Budesonide. Pharmaceutics 12.

65. Mosharraf M, Nyström C (1995) The effect of particle size and shape on the surface specific dissolution rate of microsized practically insoluble drugs. Int J Phar 122: 35–47.

66. Rohrschneider M (2012) Correlation of ICS in vitro dissolution and pulmonary absorption (Thesis). Düsseldorf, Germany: Heinrich Heine University Düsseldorf.

67. Schilling U (2017) The role of in vitro and pharmacokinetics studies in the bioequivalence assessment of inhaled and intranasal corticosteroids (Thesis): University of Florida.

68. Kumar A, Terakosolphan W, Hassoun M, Vandera KK, Novicky A, et al. (2017) A Biocompatible Synthetic Lung Fluid Based on Human Respiratory Tract Lining Fluid Composition. Pharm Res 34: 2454–65.

69. Meindl C, Stranzinger S, Dzidic N, Salar-Behzadi S, Mohr S, et al. (2015) Permeation of Therapeutic Drugs in Different Formulations across the Airway Epithelium In Vitro. PLoS One 10: e0135690.

70. Lee MK, Yoo JW, Lin H, Kim YS, Kim DD, et al. (2005) Air-liquid interface culture of serially passaged human nasal epithelial cell monolayer for in vitro drug transport studies. Drug Deliv 12: 305–11.

71. Crowe A, Tan AM (2012) Oral and inhaled corticosteroids: differences in P-glycoprotein (ABCB1) mediated efflux. Toxicol Appl Pharmacol 260: 294–302.

72. Siebert TA, Rugonyi S (2008) Influence of liquid-layer thickness on pulmonary surfactant spreading and collapse. Biophys J 95: 4549–59.

73. Pattle RE (1955) Properties, function and origin of the alveolar lining layer. Nature 175: 1125–6.

74. Wanner A (1977) Clinical aspects of mucociliary transport. Am Rev Respir Dis 116: 73–125.

75. Widdicombe J (1997) Airway and alveolar permeability and surface liquid thickness: theory. J Appl Physiol (1985) 82: 3–12.

76. Kondo T, Hibino M, Tanigaki T, Cassan SM, Tajiri S, et al. (2017) Appropriate use of a dry powder inhaler based on inhalation flow pattern. J Pharm Health Care Sci 3: 5.

77. Tamura G, Sakae H, Fujino S (2012) In vitro evaluation of dry powder inhaler devices of corticosteroid preparations. Allergol Int 61: 149–54.

78. Bustamante-Marin XM, Ostrowski LE (2017) Cilia and Mucociliary Clearance. Cold Spring Harb Perspect Biol 9.

79. Hofmann W, Asgharian B (2003) The effect of lung structure on mucociliary clearance and particle retention in human and rat lungs. Toxicol Sci 73: 448–56.

80. Wang J, Flanagan DR (1999) General solution for diffusion-controlled dissolution of spherical particles. 1. Theory. J Pharm Sci 88: 731–8.

81. Wang J, Flanagan DR (2002) General solution for diffusion-controlled dissolution of spherical particles. 2. Evaluation of experimental data. J Pharm Sci 91: 534–42.

82. Kannan R, Przekwas A (2020) A multiscale absorption and transit model for oral drug delivery: Formulation and applications during fasting conditions. International Journal for Numerical Methods in Biomedical Engineering 36: e3317.

83. Cheng YS (2014) Mechanisms of pharmaceutical aerosol deposition in the respiratory tract. AAPS PharmSciTech 15: 630–40.

84. Olsson B, Kassinos SC (2021) On the Validation of Generational Lung Deposition Computer Models Using Planar Scintigraphic Images: The Case of Mimetikos Preludium. J Aerosol Med Pulm Drug Deliv 34: 115–23.

85. Harrison TW, Tattersfield AE (2003) Plasma concentrations of fluticasone propionate and budesonide following inhalation from dry powder inhalers by healthy and asthmatic subjects. Thorax 58: 258–60.

86. Mollmann H, Wagner M, Krishnaswami S, Dimova H, Tang Y, et al. (2001) Single-dose and steady-state pharmacokinetic and pharmacodynamic evaluation of therapeutically clinically equivalent doses of inhaled fluticasone propionate and budesonide, given as Diskus or Turbohaler dry-powder inhalers to healthy subjects. J Clin Pharmacol 41: 1329–38.

87. Mortimer KJ, Harrison TW, Tang Y, Wu K, Lewis S, et al. (2006) Plasma concentrations of inhaled corticosteroids in relation to airflow obstruction in asthma. Br J Clin Pharmacol 62: 412–19.

88. Thorsson L, Edsbacker S, Conradson TB (1994) Lung deposition of budesonide from Turbuhaler is twice that from a pressurized metered-dose inhaler P-MDI. Eur Respir J 7: 1839–44.

89. Thorsson L, Edsbacker S, Kallen A, Lofdahl CG (2001) Pharmacokinetics and systemic activity of fluticasone via Diskus and pMDI, and of budesonide via Turbuhaler. Br J Clin Pharmacol 52: 529–38.

90. Mortimer KJ, Tattersfield AE, Tang Y, Wu K, Lewis S, et al. (2007) Plasma concentrations of fluticasone propionate and budesonide following inhalation: effect of induced bronchoconstriction. Br J Clin Pharmacol 64: 439–44.

91. Vutikullird AB, Gillespie M, Song S, Steinfeld J (2016) Pharmacokinetics, Safety, and Tolerability of a New Fluticasone Propionate Multidose Dry Powder Inhaler Compared With Fluticasone Propionate Diskus((R)) in Healthy Adults. J Aerosol Med Pulm Drug Deliv 29: 207–14.

92. Gillespie M, Song S, Steinfeld J (2015) Pharmacokinetics of fluticasone propionate multidose, inhalation-driven, novel, dry powder inhaler versus a prevailing dry powder inhaler and a metered-dose inhaler. Allergy Asthma Proc 36: 365–71.

93. van den Brink KI, Boorsma M, Staal-van den Brekel AJ, Edsbacker S, Wouters EF, et al. (2008) Evidence of the in vivo esterification of budesonide in human airways. Br J Clin Pharmacol 66: 27–35.

94. Movia D, Prina-Mello A (2020) Preclinical Development of Orally Inhaled Drugs (OIDs)- Are Animal Models Predictive or Shall We Move Towards In Vitro Non-Animal Models? Animals (Basel) 10.

95. Asgharian B, Hofmann W, Bergmann R (2001) Particle Deposition in a Multiple-Path Model of the Human Lung. Aerosol Science and Technology 34: 332–9.

96. Phalen RF, Cuddihy RG, Fisher GL, Moss OR, Schlesinger RB, et al. (1991) Main Features of the Proposed NCRP Respiratory Tract Model. Radiation Protection Dosimetry 38: 179–84.

97. Robinson RJ, Snyder P, Oldham MJ (2008) Comparison of analytical and numerical particle deposition using commercial CFD packages: impaction and sedimentation. Inhal Toxicol 20: 485–97.

98. Thorsson L, Dahlstrom K, Edsbacker S, Kallen A, Paulson J, et al. (1997) Pharmacokinetics and systemic effects of inhaled fluticasone propionate in healthy subjects. Br J Clin Pharmacol 43: 155–61.

99. BurM, Rothen-Rutishauser B, Huwer H, Lehr CM (2009) A novel cell compatible impingement system to study in vitro drug absorption from dry powder aerosol formulations. Eur J Pharm Biopharm 72: 350–57.

100. Haghi M, Ong HX, Traini D, Young P (2014) Across the pulmonary epithelial barrier: Integration of physicochemical properties and human cell models to study pulmonary drug formulations. Pharmacol Ther 144: 235–52.

101. Haghi M, Traini D, Postma DS, Bebawy M, Young PM (2013) Fluticasone uptake across Calu-3 cells is mediated by salmeterol when deposited as a combination powder inhaler: Salmeterol-mediated fluticasone uptake. Respirology 18: 1197–1201.

102. Hein S, Bur M, Schaefer UF, Lehr CM (2011) A new Pharmaceutical Aerosol Deposition Device on Cell Cultures (PADDOCC) to evaluate pulmonary drug absorption for metered dose dry powder formulations. Eur J Pharm Biopharm 77: 132–8.

103. Himstedt A, Braun C, Wicha SG, Borghardt JM (2020) Towards a Quantitative Mechanistic Understanding of Localized Pulmonary Tissue Retention-A Combined In Vivo/In Silico Approach Based on Four Model Drugs. Pharmaceutics 12.

104. Newman B, Witzmann K (2020) Addressing the Regulatory and Scientific Challenges with Generic Orally Inhaled Drug Products. Pharmaceut Med 34: 93–102.

105. Forbes B, Asgharian B, Dailey LA, Ferguson D, Gerde P, et al. (2011) Challenges in inhaled product development and opportunities for open innovation. Adv Drug Deliv Rev 63: 69–87.

106. Kirby AC, Coles MC, Kaye PM (2009) Alveolar macrophages transport pathogens to lung draining lymph nodes. J Immunol 183: 1983–89.

107. Rubin K, Ewing P, Backstrom E, Abrahamsson A, Bonn B, et al. (2020) Pulmonary Metabolism of Substrates for Key Drug-Metabolizing Enzymes by Human Alveolar Type II Cells, Human and Rat Lung Microsomes, and the Isolated Perfused Rat Lung Model. Pharmaceutics 12.

