## Supporting Information for "Predicting systemic and pulmonary tissue barrier concentration of orally inhaled drug products"

S1 Appendix

Narender Singh, Ravi Kannan, Ryan Arey, Ross Walenga, Andrew Babiskin, and Andrzej Przekwas

### Contents

**1 Deposition computations**
**2 Transport and deposition equation, for the aerosol species transport**
**3 Mesh refinement analysis**

### 1. Deposition computations

The biggest assumption for deposition computations is that the individual monodispersed particles do not affect each other. This assumption allows us to instantiate multiple invocations of the species transport module, corresponding to different particle sizes. Another major assumption is that complex 3D phenomena like flow-recirculation should not overtly affect the deposition process. This way, we can plug analytical deposition expressions in the Euler-Euler Q3D module, in each computational cell.

Since the transport is calculated using a species transport equation, the species value is set to zero for the cell averaged concentrations. The inlet concentration $C_{INLET}$ is set to 1.0, during the actuation period. It is set to zero after the actuation period. Thus, the total mass inhaled (for that mono-dispersed particle class) is $M_{INLET} = \int_0^{T_{inhale}} C_{INLET}.\ A_{INLET}\ U_{INLET} dt$, where $U_{INLET}$ is the aerosol inlet velocity.

The deposition process occurs on the Q3D walls. The deposition fraction in each computational cell is given by: $\frac{\int_0^{\infty} C_{ENTRY}.\ U.\pi R^2\ P_{TOTAL} dt}{\int_0^{T_{inhale}} C_{INLET}.\ A_{INLET}\ U_{INLET} dt}$, where $P_{TOTAL}$ is the overall deposition probability, U is the mean particle transport velocity in that computational cell, R is the radius of the airway segment and $C_{ENTRY}$ is the concentration at which the aerosol enters the computational cell.

The first step is to compute the probability of deposition in each computational Q3D cell. There are three methods of deposition that can occur. They are described below:

The probability of the diffusion deposition in each Q3D mesh segment, $P_{DIFF}$, is obtained from the analytical expressions for the diffusion deposition in a tube by multiple research programs [1-3]. As noted by Ingham et al [3], several terms of the infinite series equation are needed to compute the end concentration. By limiting the expression to the first 5 terms, we obtain a good approximation for the probability of the diffusional deposition:

$$P_{DIFF} = 0.819\, e^{-7.315x} + 0.0976\, e^{-44.61x} + 0.0325\, e^{-114x} + 0.0509 e^{-79.31x^{2/3}} \quad (1)$$

where,

$$x = \frac{(Length/U_{Mean})}{2R_0^2/D} \quad (2)$$

L is the length of the airway segment, U is the mean particle transport velocity in that computational cell, R is the radius of the airway segment, and D is the diffusion coefficient of particles in air given by: $D = \frac{k_B T C_{corr}}{6\pi\mu r_p}$. $k_B$ is the Boltzmann constant, T is the body temperature, $r_p$ is the microparticle radius, μ is the air viscosity. Ccorr is the Cunningham slip correction factor [4], and is given by: $C_{corr} = 1 + \frac{2\lambda}{2r_p}(A_1 + A_2\, exp\left({-A_3 2r_p}/{\lambda}\right)$. For air, $A_1$=1.257, $A_2$ = 0.400, $A_3$ = 0.55 [5]. $\lambda$ (80 nm) is the mean free path. For $r_p$=1.1 µm, $C_{corr}$ = 1.086 because $\lambda$ is much smaller than the microparticle radius $r_p$.

The probability of the impaction in a non-bifurcating tube can be obtained from the semi-empirical formulation of Weiden et al [6] and Pui et al [7]: $P_{IMPACT} = 1 - (1 + (\frac{Stk}{0.171})^{0.452\frac{Stk}{0.171}+2.242})^{\frac{-2\theta}{\pi}}$ where $\theta$ is the angle of curvature of the bend and $Stk$ is the local Stokes number.
The probability of the deposition due to the acceleration due to the gravity (i.e., sedimentation) is given by [6]: $P_{SEDI} = 1 - exp\left(-\frac{4gC_{corr}\rho {r_p}^2 L \cos\varphi}{9\pi\mu RU}\right)$ where, $\rho$ is the particle density, g is the gravity, $\phi$ is the inclination angle of the airway segment relative to the gravity (0 for the horizontal tube) and $C_{corr}$ is the Cunningham slip correction factor. The efficiency $P_{IMPACT}$ due to impaction deposition, in the case of bifurcating airways is given by [8]:

$$P_{INPACT} = 1 - \frac{2}{\pi}\cos^{-1}(St.\theta) + \frac{1}{\pi}\sin(2\cos^{-1}(St.\theta)); St.\theta < 1$$
$$P_{INPACT} = 1; St.\theta \geq 1 \quad (3)$$

where θ is the branching angle of the airway generation and *St* is the Stokes' number. The overall deposition probability is given by $P_{TOTAL} = 1 - (1 - P_{DIFF})(1 - P_{IMPACT})(1 - P_{SEDI})$. This loss term is implemented in the species transport equation, in CFDRC's finite volume method CoBi. Additional details on the species or flow transport can be obtained from previous CoBi publications [9-12].

### 2.Transport and deposition equation, for the aerosol species transport

In every computational Q3D cell, the following equations relate the entry, cell average and exit concentration states:

$$\frac{C_{in} + C_{OUT}}{2} = \bar{C}; C_{OUT} = (1 - P)C_{in}$$

The rate of mass lost, due to the deposition in the Q3D cell is given by:

$\dot{M}_{LOSS} = Q(C_{in} - C_{OUT}) = QC_{in}(P) = \frac{2PQ\bar{C}}{2-P}$,
Where Q is the flowrate. Plugging this into the species transport equation and integrating over the Q3D cell results in the following equation:

$$Vol.\frac{\partial \bar{C}}{\partial t} + \oint (\nabla(u_i C) - D_i \nabla^2(C))dV = -\frac{2PQ\bar{C}}{2-P}$$

**3.Mesh refinement analysis**

The simulations performed using the whole lung Q3D model are mesh independent. Some metrics are provided here to support mesh independence. The normalized error between the fine (~187K cells) and the medium meshes (~95 K cells) are provided.

| **Region** | **Normalized error between the two meshes** |
|---|---|
| Respiratory region | 4% |
| TB region | 3% |

**References**


1. Goldstein S. Modern developments in fluid dynamics. *The Oxford Engineering Science Series*. 1938.
2. Gormley PG and Kennedy M. Proc. R. Ir. Acad. 52, 163. 1949.
3. Ingham DB. Diffusion of aerosols, from a stream moving through a cylindrical tube. *Aerosol Science*. Vol 6, pp 125-32. 1975.
4. Cunningham, E., On the velocity of steady fall of spherical particles through fluid medium. *Proc. Roy. Soc. A* 83: 357-65, 1910
5. Davies, CN. Definitive equations for the fluid resistance of sphere. *Proc. Phys. Soc. London*, 57:259-70, 1945.
6. Weiden V, Drewnick F and Borrmann S. Particle Loss Calculator – a new software tool for the assessment of the performance of aerosol inlet systems. *Atmos. Meas. Tech*., 2, 479-94, 2009.
7. Pui D, Romay-Novas D and Liu B. Experimental Study of Particle Deposition in Bends of Circular Cross Section. *Aerosol Science and Technology*. 7:3, 301-315. 1987
8. Anjilvel S, Asgharian B. A multiple-path model of particle deposition in the rat lung. *Fundam Appl Toxicol*. 1995 Nov;28(1):41-50.
9. Kannan R, Przekwas A. A near-infrared spectroscopy computational model for cerebral hemodynamics. *Int J Numer Method Biomed Eng*. 2012 Nov;28(11):1093-106.
10. Kannan R, Przekwas A. A computational model to detect and quantify a primary blast lung injury using near-infrared optical tomography. *Int J Numer Method Biomed Eng*. 2012 Jan;27(11):13-28.
11. Kannan RR, Singh N, Przekwas A. A compartment-quasi-3D multiscale approach for drug absorption, transport, and retention in the human lungs. *Int J Numer Method Biomed Eng*. 2018 May;34(5):e2955.
12. Kannan RR, Singh N, Przekwas A. A Quasi-3D compartmental multi-scale approach to detect and quantify diseased regional lung constriction using spirometry data. *Int J Numer Method Biomed Eng*. 2018 May;34(5):e2973.
